# Thymic T_REG_-derived T follicular regulatory cells prevent catastrophic humoral autoimmunity in response to TLR7-driven inflammation

**DOI:** 10.64898/2026.06.28.735092

**Authors:** Layla Pohl, Lasse F. Voss, Gudrun Winther, Sara Simone Jensen, Kenneth Green, Yamira C. L. Weber, Julia Karen Demtröder, Cecilia Fahlquist-Hagert, Thomas R. Wittenborn, Chaido Sirinian, Mathias Krogh Pedersen, Johanne Vium-Heinesen, Kristian Savstrup Kastberg, Sofie V. Fonager, Johannes Skraep, Ali Shahrokhtash, Amanda Juul Howarth, Lisbeth Jensen, Alain Pulfer, Yonglun Luo, Duncan S. Sutherland, Anastasios D. Papanastasiou, Santiago F. Gonzalez, Anne Troldborg, Johan Palmfeldt, Lin Lin, Søren E. Degn

## Abstract

T follicular regulatory cells (T_FR_) are a follicle-resident subset of regulatory T cells (T_REG_) that limit germinal center (GC) responses and enforce humoral tolerance. Although T_FR_ are known to focus antibody responses to foreign antigens, how GC-specific regulation is maintained during inflammation remains unclear. Here, TLR7-driven inflammation unmasked a critical, non-redundant role for T_FR_ in preserving immune tolerance. Selective loss of T_FR_ caused severe autoimmunity and increased mortality, driven by inflammation-induced expansion of autoreactive B and T cell clones and epitope spreading. Autoreactivity resolved upon cessation of inflammation in T_FR_-sufficient mice but persisted in their absence. Mechanistically, T_FR_ limited the establishment and expansion of spontaneous GCs in response to inflammation, and responding T_FR_ displayed transcriptional, phenotypic, and clonal features of thymic T_REG_. This suggests that whereas T_FH_ upregulate FoxP3 to shut down end-stage GCs, it is thymic T_REG_-derived T_FR_ that safeguard GC integrity under inflammatory stress to prevent lethal autoimmunity.

## INTRODUCTION

Control of immune reactivity is essential to prevent allergy and autoimmunity. B cells are subject to multiple tolerance checkpoints during development, including negative selection and receptor editing in the bone marrow ^1^ and follicular exclusion at the transitional stage ^2^. However, upon antigen encounter, B cells can re-enter diversification programs through somatic hypermutation, particularly within germinal centers (GCs) ^3,4,5^, creating a renewed risk of autoreactivity ^6^. This risk is partially constrained by the requirement for cognate T cell help at each cycle of GC selection ^7^, yet tolerance can be breached at the level of T follicular helper cells (T_FH_) ^8,9^, and progressive epitope spreading is a hallmark of autoimmune disease ^10,11,12^. An additional layer of control is provided by a specialized subset of regulatory T cells (T_REG_), termed T follicular regulatory cells (T_FR_). ^13,14,15^. During foreign-antigen-driven GC responses, T_FR_ focus humoral immunity on the eliciting antigen and limit the emergence of auto– and allergen-reactive clones ^16,17,18^.

Whereas some studies have suggested that the T cell receptor (TCR) repertoires of T_FH_ and T_FR_ are distinct in foreign antigen-driven immune responses, others have reported that T_FR_ can be specific for the immunizing antigen and may derive from naïve T cells ^19,20^. More recently, T_FH_ in end-stage foreign antigen-driven GCs were shown to upregulate FoxP3 to acquire regulatory features ^21^. A similar phenomenon has been described in human tonsils ^22^. This has raised fundamental questions about the contributions of bona fide T_REG_-derived T_FR_ versus transient FoxP3-expressing T_FH_ in control of GC reactions ^23^.

Spontaneous GCs are also a common feature of autoimmune disease ^24^, yet the relative roles of such T_REG_– vs. T_FH_-derived T_FR_ in autoantigen-driven GCs have not been explored. Notably, in end-stage foreign antigen-driven GCs, induction of FoxP3 and acquisition of regulatory capacity in T_FH_ was linked to limiting antigen availability ^21^. By contrast, in many autoimmune diseases, the autoantigens persist and cannot be cleared, raising the question whether T_FH_– and T_REG_-derived T_FR_ are differentially regulated in the foreign antigen vs. autoimmune setting.

Chronic inflammation is linked to the development of autoimmunity ^25^. Systemic lupus erythematosus (SLE) is characterized by both extrafollicular responses and autoreactive GCs, resulting in expanded plasma cell (PC) compartments, and the production of anti-nuclear autoantibodies ^24,26,27^. Patients often experience disease flares preceded by increased inflammatory markers ^28,29,30^, and accumulating evidence implicates B cells recognizing ribonuclear antigens as key drivers of SLE pathogenesis, fueled by TLR7-driven inflammatory signaling ^31,32,33,34,35^.

Here, we leveraged a murine model of SLE driven by sterile inflammation, induced by repeated epicutaneous application of the imidazoquinoline compound R848 (resiquimod) ^36^, a TLR7 agonist ^37^. Whereas T_REG_ are required throughout the lifespan of mice to prevent catastrophic autoimmunity ^38,39^, genetic blockade or depletion of T_FR_ has previously been associated with comparatively subtle phenotypes ^13,17^. Unexpectedly, we found that TLR7-driven inflammation precipitated severe autoimmunity with significant mortality in T_FR_-deficient mice. Under milder inflammatory conditions, T_FR_ deficiency instead led to exaggerated GC responses and epitope spreading. Notably, cessation of inflammatory stimulation resulted in resolution of autoreactivity in wild-type mice, whereas autoimmunity persisted in a subset of T_FR_-deficient animals. Mechanistically, FoxP3-expressing cells emerged early during spontaneous GC formation and inversely correlated with GC expansion but did not correlate with GC contraction, indicating that these were T_REG_-derived T_FR_, not phenoconverted T_FH_. Moreover, they expressed the lineage marker Helios ^40^ and exhibited transcriptional and clonal features overlapping with thymic T_REG_, suggesting that thymic T_REG_-derived T_FR_ are key regulators of early, inflammation-driven GC responses. We further show that circulating T_FR_ frequencies correlate with disease activity in patients with SLE, suggesting that T_FR_ expand during disease flares to exert GC control.

## RESULTS

### Sterile inflammation causes severe autoimmunity in T_FR_-deficient mice

Sterile inflammation by repeated epicutaneous application of R848 causes a phenotype in mice that recapitulates hallmarks of SLE ^36^. T_FR_ require both the transcription factors FoxP3 and Bcl6, allowing us to test the role of T_FR_ in restricting spontaneous GC responses to inflammation by combining R848 treatment with previously established ^41^ conditional FoxP3-Cre-driven deletion of Bcl6 (FoxP3-Cre;Bcl6^flx/flx^) (Fig. 1A). To corroborate the genetic strategy, we performed spatial analyses of spleen sections, confirming that T_FR_ were present in GCs of wild-type C57BL6/J (B6) mice but absent in FoxP3-Cre;Bcl6^flx/flx^ mice (Fig. S1A and B), henceforth referred to as T_FR_-sufficient and deficient mice, respectively.

**Figure 1.**
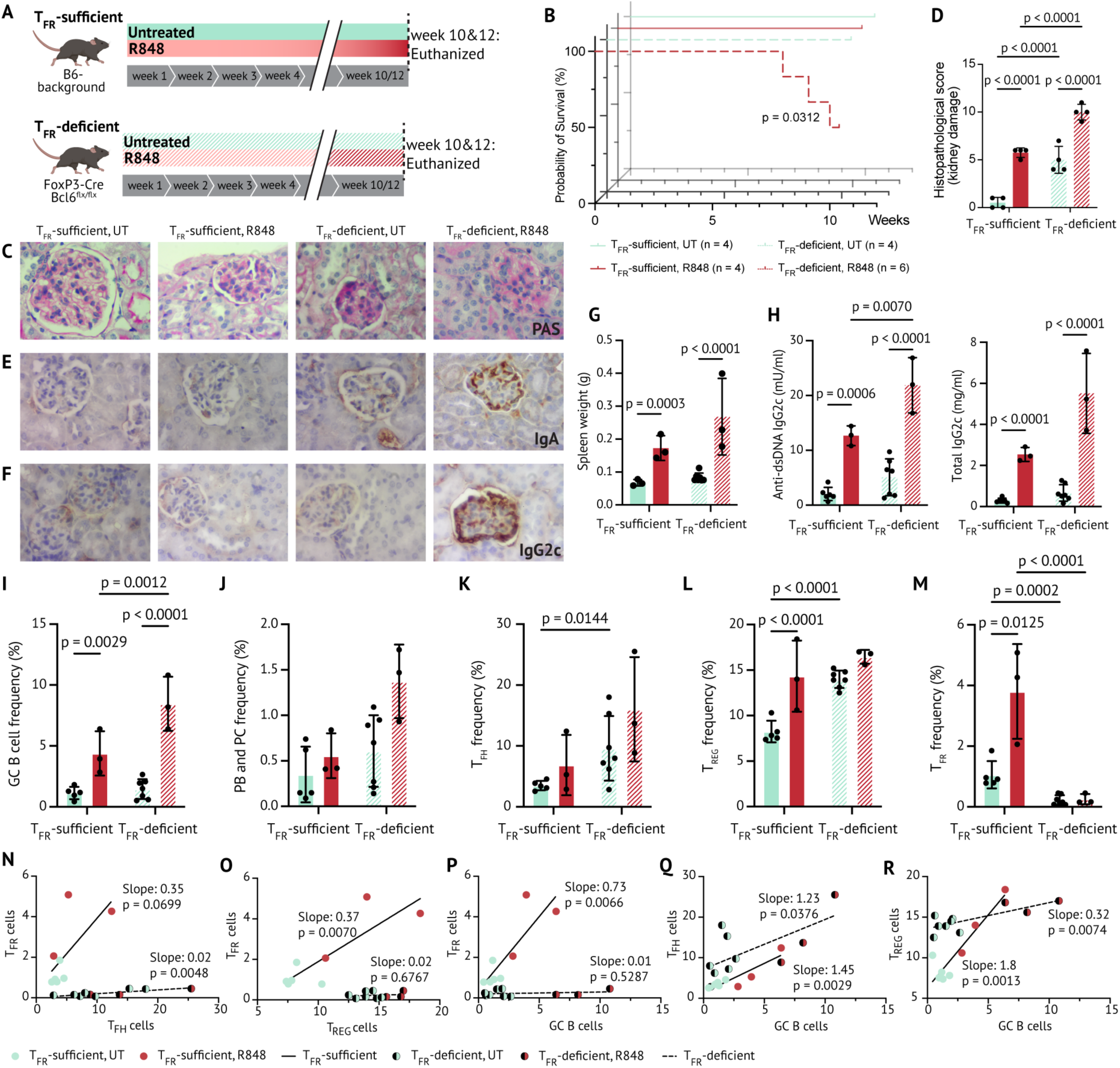
Sterile inflammation causes severe autoimmunity in T_FR_-deficient mice. (A) Overview of experimental setup for high-dose (for 10 weeks, panels B-F) and low-dose (for 12 weeks, panels G-R) R848 treatment regimens. (B) Kaplan–Meier survival curves for untreated (UT) and R848-treated T_FR_-suLicient and deficient mice. Groups were compared using the log-rank test. Starting n is given in parenthesis in the legend. Tissue samples could not be recovered from two of the deceased mice, resulting in n = 4 in all groups in subsequent analysis presented in panels C-F. (C) Representative PAS-stained kidney sections (week 10, 40x magnification). (D) Grading of glomerular lesions of T_FR_-suLicient and deficient mice after 10 weeks of R848 treatment or no treatment (n=4 mice per group). Scores were calculated from at least 50 glomeruli per mouse, and the graph shows values for individual mice with mean±SD. P-values were computed using two-way ANOVA with uncorrected Fisher’s LSD multiple comparisons test. (E and F) Representative kidney sections stained for IgA or IgG2c deposition by immunohistochemistry (week 10, 40x magnification). (G) Spleen weights of T_FR_-suLicient and deficient mice after 12 weeks of R848 treatment or no treatment. Individual datapoints and mean±SD for n=3-7 mice per group, pooled from two cohorts. P-values were computed on log-transformed data using two-way ANOVA with uncorrected Fisher’s LSD multiple comparisons test. (H) Serum anti-dsDNA IgG2c (left panel) and total IgG2c (right panel). Individual datapoints and mean±SD for n=3-7 mice per group, pooled from two cohorts. P-values were computed using two-way ANOVA with uncorrected Fisher’s LSD multiple comparisons test. For total IgG2c, data were log-transformed to meet normality requirements for statistical analysis. (I-M) Respectively, GC B, PB and PC, T_FH_, T_REG_, and T_FR_ frequencies in spleen, determined by flow cytometry (see gating strategy in Figure S1). Individual datapoints and mean±SD for n=3-7 mice per group, pooled from two cohorts. P-values were computed using two-way ANOVA with uncorrected Fisher’s LSD multiple comparisons test. For PB and PC, T_FH_, T_REG_ and T_FH_, data were log-transformed to meet normality requirements for statistical analysis. (N) Scatterplot of T_FR_ versus T_FH_ frequencies for T_FR_-suLicient (half dots) and deficient (full dots) mice. Colors indicate R848 treatment (red) vs. no treatment (teal). Lines represent linear regression fits for each group, with slope and p-value for test of significant deviation from zero shown below the plots. (O) Scatterplot of T_FR_ frequency versus T_REG_ frequency, otherwise as N. (P) Scatterplot of T_FR_ frequency versus GC B frequency, otherwise as N. (Q) Scatterplot of T_FH_ frequency versus GC B frequency, otherwise as N. (R) Scatterplot of T_REG_ frequency versus GC B frequency, otherwise as N.

**Figure S1.**
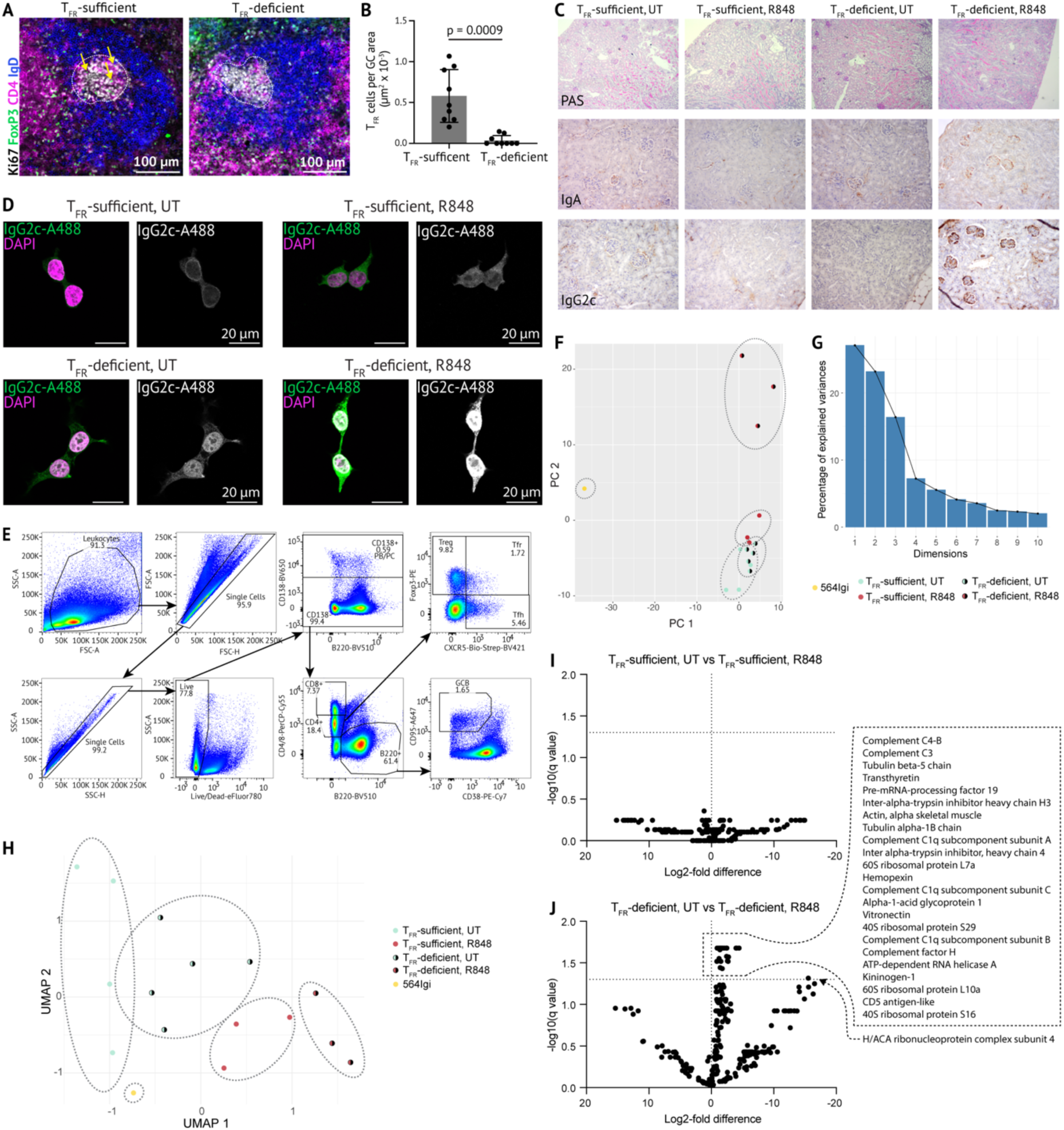
Autoreactivity and epitope spreading in T_FR_-deficient mice experiencing inflammation. (A) Representative spleen sections of T_FR_-suLicient and T_FR_-deficient mice, stained for Ki67 (grey), FoxP3 (green), CD4 (magenta), and IgD (blue). FoxP3^+^ cells were counted in IgD exclusion zones and normalized by measured GC area (IgD exclusion zone, marked by a white dashed line). Yellow arrows mark example FoxP3^+^ cells in a GC from a T_FR_-suLicient mouse, whereas in a GC from a T_FR_-deficient mouse, no FoxP3^+^ cells can be seen. Scale bar: 100 µm. (B) Quantification of T_FR_ per GC area. Individual datapoints and mean±SD are shown for n=3 mice per group (based on 3 GCs per mouse). P-value was computed using an unpaired, two-tailed t-test with Welch’s correction. (C) PAS (4x magnification), IgA and IgG2c (10x magnification) immunohistochemically stained kidney sections from T_FR_-suLicient or deficient mice after 10 weeks of R848 treatment or no treatment. Representative of 4 mice per group. (D) Anti-nuclear antibody (ANA) staining of HEK293TN cells using sera from T_FR_-suLicient or deficient mice after 12 weeks of R848 treatment or no treatment. Representative of 3 mice per group. Cells were counterstained with DAPI (nuclei, magenta) and Alexa Fluor 488-conjugated anti-IgG2c antibody was used to detect autoantibody binding (green). The Alexa Fluor 488 channel is additionally shown in grayscale to highlight antibody localization. Scale bar: 20 µm. (E) Gating strategy for data presented in main Figure 1. (F) Principal Component Analysis (PCA) of mass spectrometry-identified autoantigens targeted by antibodies in sera of T_FR_-suLicient or deficient mice treated with R848 for 10 weeks or untreated, using a 564Igi mouse as outgroup. Each point represents a mouse, colored by genotype and treatment (n=3-4 mice per group, highlighted by black dashed lines). (G) Scree plot displaying the proportion of total variance explained by each principal component derived from the dataset visualized in F. (H) UMAP plot showing global diLerences among samples based on scaled protein expression from the quantitative mass spectrometry dataset in F after removal of Ig elements and keratins. Each point represents a single mouse, colored by genotype and treatment group. Black dashed lines highlight groups. (I) Volcano plots illustrating diLerential autoantigen targeting by sera of R848-treated (n=3) versus untreated (n=4) T_FR_-suLicient mice. Each point represents an individual protein, plotted by log₂-fold change versus –log_10_(q-value). The dashed line at y = 1.3 indicates the significance threshold of q = 0.05. (J) As I, but comparing autoantigen targeting by sera of R848-treated (n=3) versus untreated (n=5) T_FR_-deficient mice.

Whereas BALB/c mice display significant morbidity and mortality upon R848 treatment, C57BL/6J mice develop a milder autoimmune phenotype ^36^. However, after 8-10 weeks of R848 treatment, we were surprised to observe that several T_FR_-deficient mice stopped thriving, with some dying unexpectedly, prompting early termination due to ethical considerations (Fig. 1B). By Kaplan-Meier survival curve analysis, R848-treated T_FR_-deficient mice exhibited significantly reduced survival compared to all other groups (p = 0.0312). No mice from any other group died spontaneously, nor did any reach a humane endpoint (Fig. 1B).

To determine whether lethality was associated with autoimmune pathology, we performed histological analyses of the kidney. Periodic Acid-Schiff (PAS)-staining revealed signs of glomerulonephritis after R848 treatment in both T_FR_-sufficient and deficient mice (Fig. 1C and Fig. S1C). The glomerular lesions were characterized by minimal hypercellularity with diffuse mesangial prominence and associated glomerular capillary collapse. In addition, most glomeruli in R848-treated T_FR_-deficient mice showed global sclerosis, diffuse fibrosis, and focal hyalinosis. Histological scoring revealed a significant increase in glomerular lesions after R848 treatment in both groups and a significantly elevated score in T_FR_-deficient compared to sufficient mice irrespective of treatment status (Fig. 1D). Furthermore, marked IgA and IgG2c immune complex deposition was observed in glomeruli of T_FR_-deficient mice, consistent with immune complex-mediated nephritis as seen in SLE, with severe manifestations upon R848-treatment (Fig. 1E and F, and Fig. S1C). Taken together, this revealed a histological phenotype indicative of glomerulopathy in untreated T_FR_-deficient mice and both groups after treatment, while pathology resembling an immune-mediated disease in humans was observed in R848-treated T_FR_-deficient mice.

### T_FR_-deficient mice show increased GCs and autoantibodies in response to inflammation

To avoid mortality and further evaluate the immunophenotype, we repeated the experiment with a lower R848 dose (used in all experiments from here on), which enabled survival of T_FR_-deficient mice over the course of 12 weeks (Fig. 1A). R848-treated mice exhibited significant splenomegaly, and this was more severe in T_FR_-deficient mice (Fig. 1G). We also observed elevated IgG2c class-switched anti-double-stranded (ds)DNA antibodies and total levels of the Th1-associated isotype IgG2c in sera of R848-treated mice, and again to a greater extent in T_FR_-deficient mice (Fig. 1H). Indirect immunofluorescence assays on HEK293TN cells revealed that serum IgG2c antibodies targeted intracellular antigens, with staining observed in both nuclear and cytoplasmic compartments upon R848 treatment (Figure S1D). Reactivity was markedly increased in R848-treated T_FR_-deficient mice but also evident in untreated T_FR_-deficient mice (Fig. S1D).

Flow cytometric analysis of the spleen showed a significant increase in the frequency of GC B, plasmablasts (PBs) and plasma cells (PCs), T_FH_, as well as T_REG_ in R848-treated mice, all trending to a higher degree in T_FR_-deficient mice, with GC B cell frequencies reaching significance (Fig. 1I-L, gating strategy in Fig. S1E). Flow cytometric analysis also revealed an increase in T_FR_ frequencies in T_FR_-sufficient mice upon R848 treatment and confirmed an effective genetic T_FR_ blockade in FoxP3-Cre;Bcl6^flx/flx^ mice (Fig. 1M). The FoxP3-Cre driver has been reported to display both a low level of hypomorphism and leakiness ^42^. However, the increased baseline T_REG_ frequency in untreated FoxP3-Cre;Bcl6^flx/flx^ compared to wild-type mice (Fig. 1L) indicated sufficient FoxP3 expression levels to support global T_REG_ differentiation, and the observed increase of T_FH_ (Fig. 1K) argued against leaky recombination of the Bcl6 target allele, because T_FH_ also require intact Bcl6. Together, this indicated that T_FR_ expand to control inflammation in wild-type mice, and corroborated that T_FR_, but not T_REG_ or T_FH_, were selectively genetically blocked in FoxP3-Cre;Bcl6^flx/flx^ mice (Fig. 1K-M).

### Epitope spreading is exacerbated in the absence of T_FR_

To assess whether R848-treated and T_FR_-deficient mice exhibited increased epitope spreading, we performed an unbiased autoantigen screen. Autoantibody-bound antigens were pulled down from pre-cleared spleen lysates from naïve mice and identified by mass spectrometry. Principal component analysis (PCA) revealed that T_FR_-blocked R848-treated mice grouped separately from their untreated counterparts and both untreated and treated T_FR_-sufficient mice (Figure S1F) and were distinct from another autoimmune model (564Igi ^35^). Of note, the first two PCs accounted for around 50% of the total variation (Fig. S1G). When all parameters were considered by UMAP dimensionality reduction, both R848-treated groups segregated together, with a progressive distance from 564Igi, untreated T_FR_-blocked mice, and finally untreated T_FR_-sufficient mice (Fig. S1H). We removed Ig and keratin-derived targets to exclude the antibodies themselves and a common background, and then directly compared untreated vs. treated mice within each group (Fig. S1I and J). Whereas there were no significant hits in the T_FR_-sufficient group, we observed an underlying broadening of targets in T_FR_-deficient mice, with multiple significant hits after R848 treatment (Fig. S1J). The pattern of reactivity in R848-treated T_FR_-deficient mice was consistent with a lupus-like, GC-driven autoimmune signature, including complement autoantigens (C1q, C3, C4, complement factor H), ribosomal and RNA-binding proteins, cytoskeletal proteins, and plasma proteins commonly found in SLE immune complexes. Hence, T_FR_-deficient mice exhibited an autoantibody profile congruent with epitope spreading during R848-induced inflammation.

### Blockade of T_FR_ shifts GC B: T_FH_ correlations and promotes T_REG_ increase

To assess how T_FR_ respond to inflammation in relation to other immune cell subsets, we plotted pairwise relationships between selected populations for each genotype (Fig. 1N–R). In T_FR_-sufficient mice, the frequencies of T_FH_, T_REG_, and GC B cells correlated with T_FR_ frequencies, consistent with a coordinated expansion in response to inflammation (Fig. 1N-P). As expected, these correlations were absent in T_FR_-deficient mice since T_FR_ were lost (Fig. 1N-P). GC B and T_FH_ cells exhibited a strong positive correlation, consistent with their co-regulated expansion during the autoimmune response (Fig. 1Q), as previously reported ^43^. The slopes of the regression lines were similar between the two groups, indicating a parallel relationship, but the intercept was higher in T_FR_-deficient mice, reflecting increased T_FH_ differentiation, proliferation, and/or persistence, again supporting that inadvertent FoxP3-Cre-driven deletion of Bcl6 did not occur in this subset. Notably, T_REG_ and GC B cells also increased in concert upon treatment of T_FR_-sufficient mice, whereas this shift was reduced in T_FR_-deficient mice, which already exhibited elevated baseline T_REG_ frequencies prior to R848 treatment (Fig. 1R).

Taken together, this showed that T_FR_ contribute to regulation of the GC niche, as previously reported ^17^, and further suggested that blocking their differentiation genetically may cause their retention within the T_REG_ population or drive a compensatory T_REG_ expansion, as indicated by the elevated baseline T_REG_ frequency.

### T_FR_ restrain B and T cell activation and clonal expansion in response to inflammation

To further evaluate the enhanced epitope spreading and altered immune cell subsets in T_FR_-deficient mice, we performed single-cell RNA sequencing of spleen and inguinal lymph node (IngLN) samples after 10-12 weeks of R848 treatment (treatment schedule as in Fig. 1A, global clustering of resulting data in Fig. S2A). Due to the naturally low frequency of T_FR_ (exacerbated by their complete absence in T_FR_-deficient groups) and their subtle marker profile, it was not possible to pin-point specific changes in the T_FR_ population in this dataset. However, we could observe the impact of T_FR_ deficiency in other immune cell subsets.

In the B cell compartment, R848 treatment induced a relative reduction in naïve, immature/transitional and marginal zone (MZ) B cells, both in T_FR_-sufficient and T_FR_-deficient groups, and more pronounced in the T_FR_-deficient group (Fig. 2A-C). Conversely, we observed an expansion of activated, atypical or age-associated memory B, GC B, plasmablast and plasma cell subsets following R848 treatment, with a more marked effect in the T_FR_-deficient group (Fig. 2A-C). The R848-treated T_FR_-deficient group displayed an expansion of GC dark zone (DZ) B cells, which were mainly found to be in either S– or G2M-phase, corroborating their identity (Fig. S2B).

**Figure 2.**
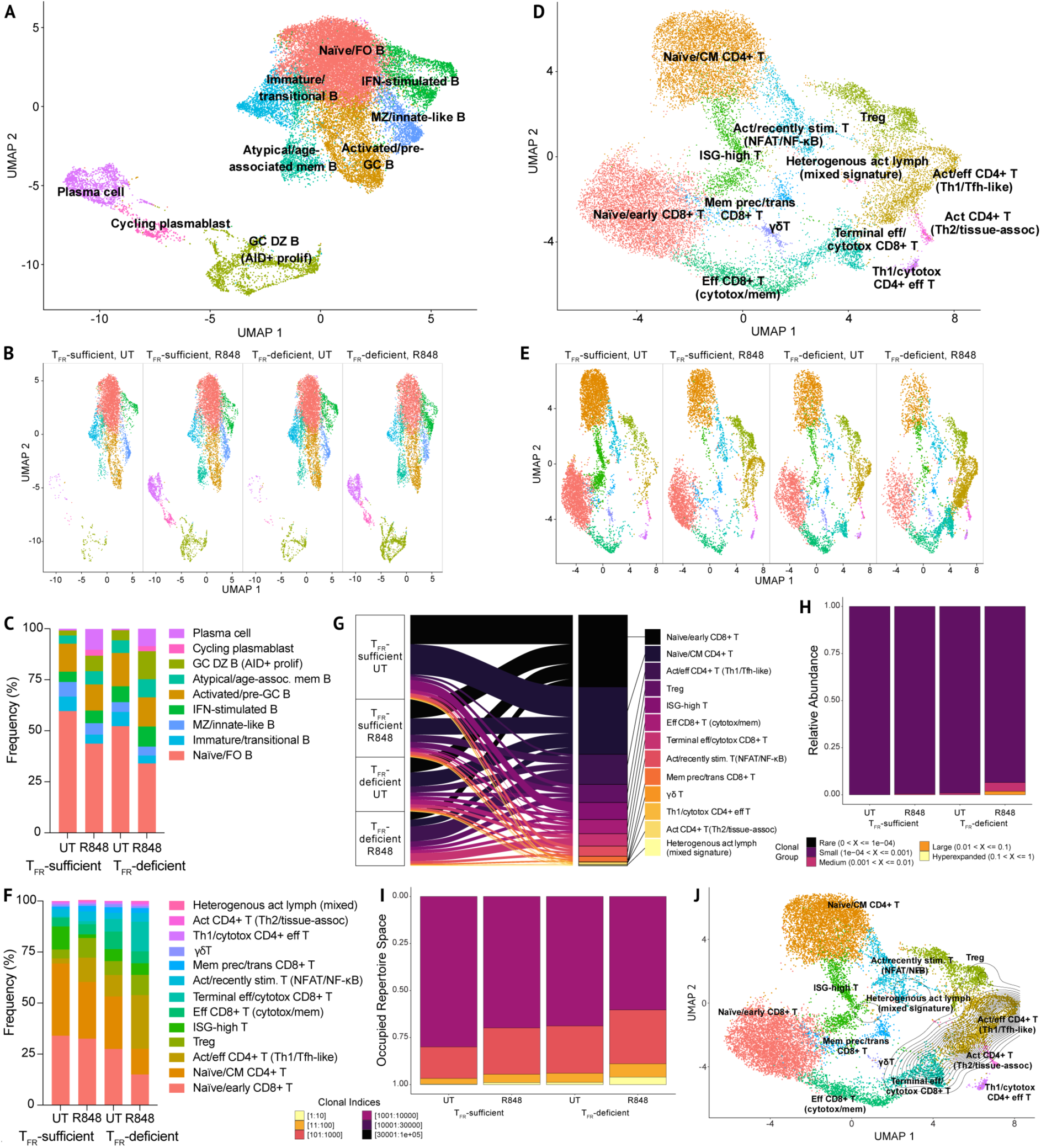
Inflammation drives increased B and T cell activation and clonal expansion in the absence of T_FR_. (A) UMAP of B cells from spleens (n = 3 mice per group, 12 total) and inguinal lymph nodes (n = 1 mouse per group, 4 total) of T_FR_-suLicient and T_FR_-deficient mice after 10-12 weeks of R848 treatment or no treatment. Each dot represents a single cell, colored by transcriptionally defined cluster. Cell types were identified from single-cell RNA-seq data (4 lanes in total from two independent experiments) using canonical marker-based clustering in Seurat. (B) UMAP of B cells split by group, otherwise as A. (C) B cell cluster frequencies. (D) UMAP of T cells, otherwise as A. (E) UMAP of T cells split by group, otherwise as A. (F) T cell cluster frequencies. (G) Alluvial plot of T cell clones, split by group and colored by cluster identities. (H) Relative abundance of expanded T cell clones by group. (I) Occupied repertoire space of T cell clones by group. (J) Contour overlay of hyperexpanded T cell clones on UMAP colored by group identity.

**Figure S2.**
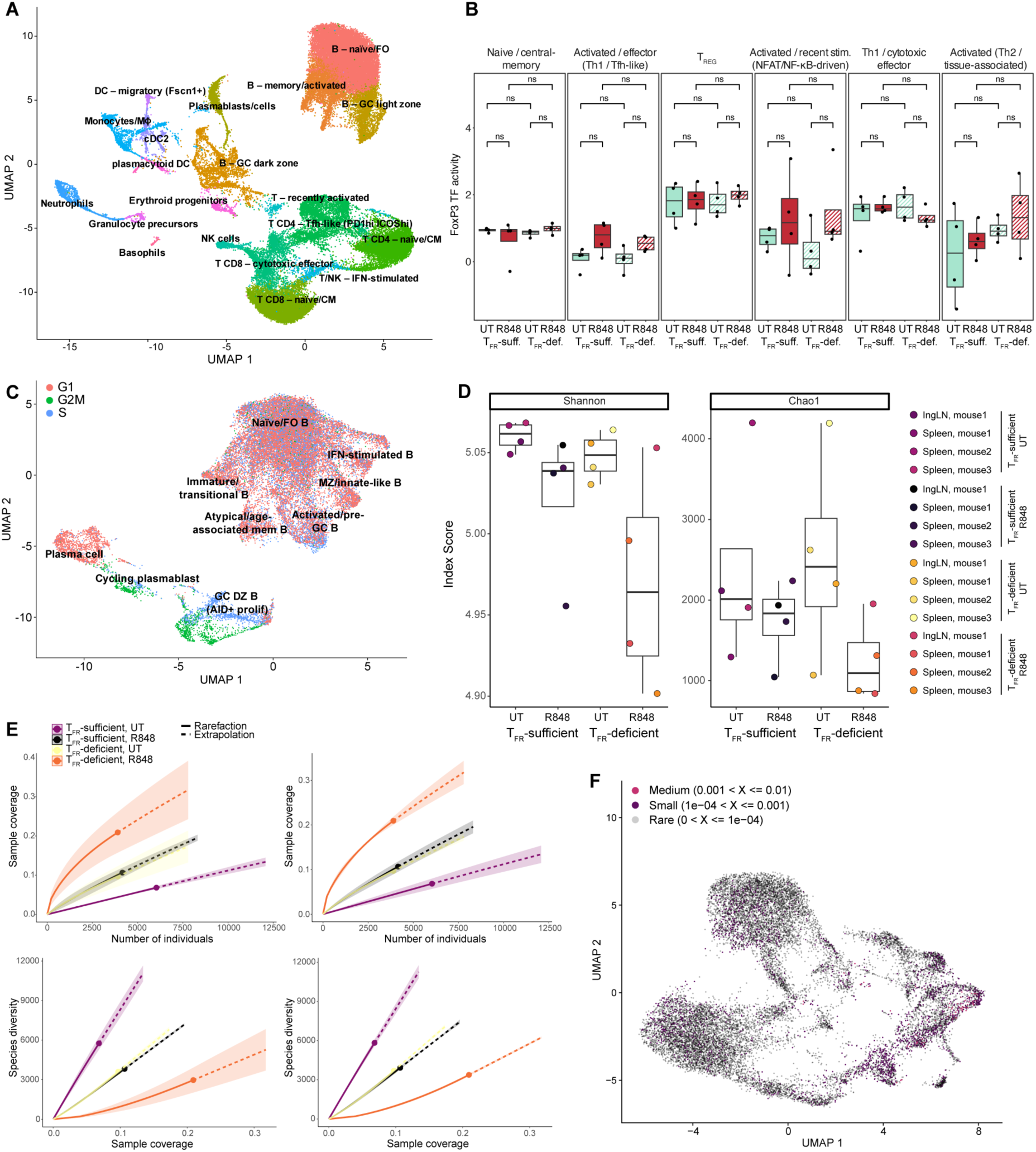
Cell and clonal dynamics in response to inflammation in the presence or absence of T_FR_. (A) UMAP of leukocytes from spleens (n = 3 mice per group, 12 total) and inguinal lymph nodes (n = 1 mouse per group, 4 total) of T_FR_-suLicient and deficient mice after 10-12 weeks of R848 treatment or no treatment. Each dot represents a single cell, colored by transcriptionally defined cluster. Cell types were identified from single-cell RNA-seq data (4 lanes in total from two independent experiments) using canonical marker-based clustering in Seurat. (B) UMAP of B cells, as in main Fig. 2A, but colored for cell cycle stage. (C) FoxP3 regulon score across the 6 CD4 T cell clusters represented in Figure 2D, based on pseudobulking and regulon transcription factor (TF) analysis using decoupleR against the DoRothEA transcription factor database. (D) Dot and box plot showing Shannon entropy (left) and Chao1 (right) estimator levels by group. Each dot represents a sample (spleen or IngLN) from a mouse (numbered), as indicated in the legend. The boxes show the interquartile ranges with medians, and whiskers extend to the most extreme values within 1.5 times the interquartile ranges. (E) T cell clonal rarefaction using Shannon diversity (q=1, lefthand plots) or species richness (q=0, righthand plots), showing either sample coverage as a function of number of individuals (clones) (top) or species diversity as a function of sample coverage (bottom). (F) UMAP of T cells, as in main Fig. 2D, but colored for clone size.

Within the T cell compartment, R848 treatment was associated with a reduction in naïve subsets and an expansion of activated effector CD4⁺ T cells exhibiting a Th1/T_FH_-like phenotype, with this effect being most prominent in the T_FR_-deficient group (Fig. 2D-F). R848 treatment induced a comparable expansion of conventional T_REG_ in both T_FR_-sufficient and T_FR_-deficient groups, indicating that the blockade of T_FR_ did not impair overall T_REG_ differentiation. Notably, T_REG_ frequencies were modestly increased in the untreated T_FR_-deficient group, consistent with either retention of cells otherwise destined for the T_FR_ compartment or a compensatory expansion of the T_REG_ pool. In addition, R848 treatment was associated with an expansion of effector and cytotoxic CD8⁺ T cells (Fig. 2D-F). To further evaluate T_REG_ functionality in the FoxP3-Cre group we performed pesudobulk FoxP3 regulon transcription factor analysis using decoupleR against the DoRothEA transcription factor database. FoxP3 transcription factor activity in the T_REG_ subset was comparable between wild-type and FoxP3-Cre;Bcl6^flx/flx^ mice (Fig. S2C), supporting that the presence of the Cre transgene did not cause significant FoxP3 hypomorphism in the model.

We next examined the T cell clonal landscape by analysis of TCR sequences. An alluvial clone plot stratified by group and cluster identity revealed that the R848-treated, and particularly the T_FR_-deficient group, contributed disproportionately to activated T cell clusters (Fig. 2G). Consistent with this, the T_FR_-deficient R848-treated group displayed an increased frequency of medium and large expanded clones (Fig. 2H), accompanied by a skewed distribution of clonal indices in which fewer clones occupied a larger fraction of the repertoire (Fig. 2I). Accordingly, Shannon entropy and Chao1 diversity metrics were significantly reduced in the treated groups, with the most pronounced reduction observed in the absence of T_FR_ (Fig. S2D). These differences were not attributable to reduced sampling depth resulting from relative B cell expansion, as clonal rarefaction analyses based on Shannon diversity and species richness demonstrated higher clonal diversity in the untreated T_FR_-sufficient group at equivalent sampling depths (Fig. S2E). Mapping clone sizes onto transcriptional clusters further showed that expanded clones preferentially localized to T cell clusters that were hyperexpanded in the T_FR_-deficient group (Fig. S2F and Fig. 2J). Together, these data indicated that T_FR_ are required to restrict inflammation-driven B and T cell activation and restrain GC clonal expansion.

### T_FR_ deficiency impairs contraction of the autoreactive response when inflammation resolves

We next examined the consequences of withdrawing the inflammatory stimulus during an established immune response. T_FR_-sufficient mice were treated with R848 for 4 weeks, after which treatment was discontinued (Fig. 3A). Subsets of mice were analyzed at baseline (week 0), during treatment (weeks 2 and 3), immediately after the last treatment (week 5), and 1, 2, or 6 weeks after treatment stop (weeks 6, 7, and 11, respectively). All time points were compared to baseline (week 0). We observed a gradual and significant increase in spleen to bodyweight ratio upon R848 treatment, exceeding a doubling after 4 weeks (Fig. 3B). This was followed by a progressive decrease in spleen to bodyweight ratio after treatment cessation (week 6 and 7), dropping back down to baseline by week 11, suggesting a complete reversion of the phenotype (Fig. 3B). Serum anti-dsDNA IgG2c levels increased similarly during and immediately following treatment, then dropped to baseline by week 11, indicating a transient antibody response (Fig. 3C). Together, this suggested that in the presence of T_FR_, the inflammatory stimulus induced only a transient autoreactive response.

**Figure 3.**
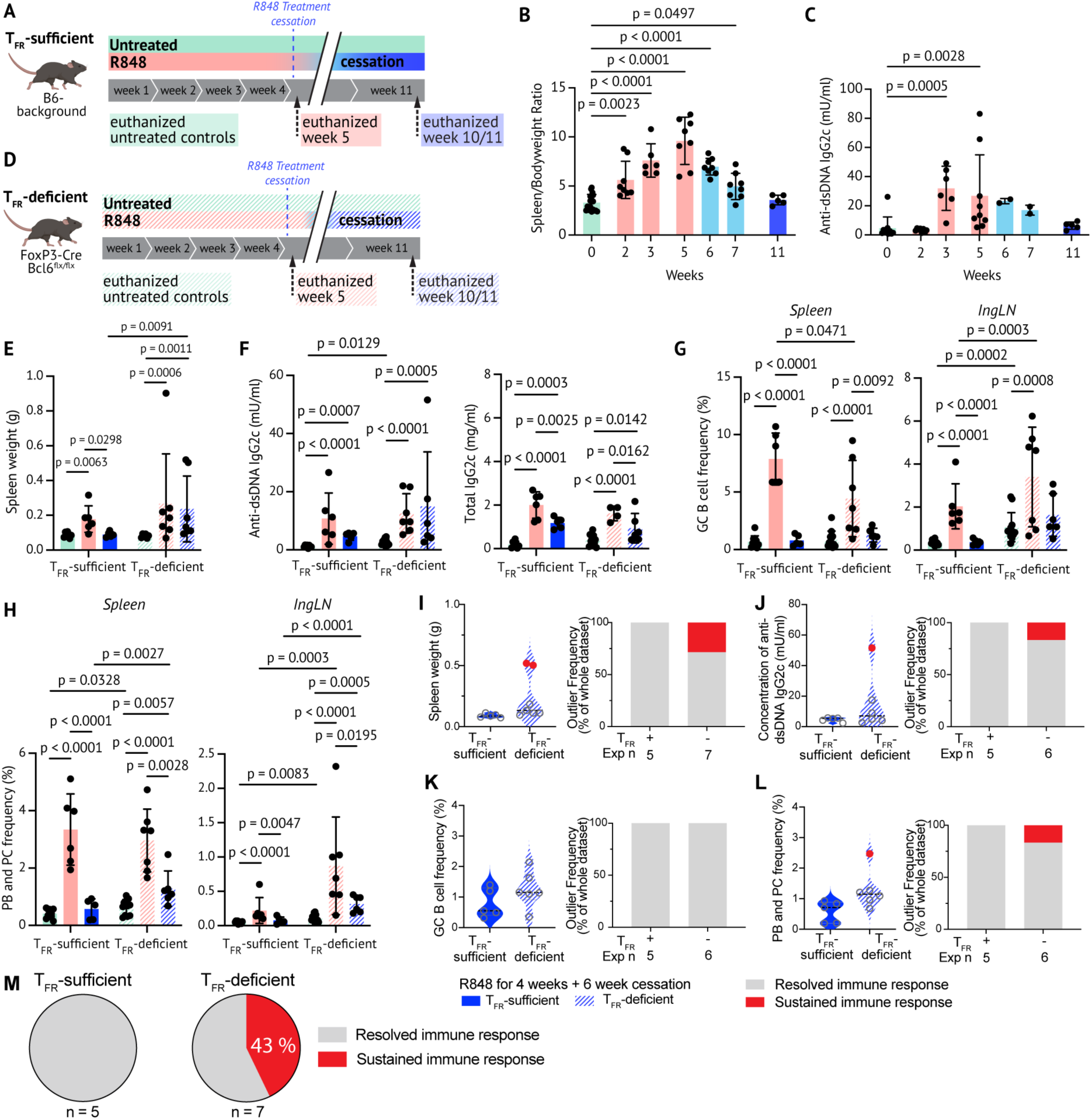
Resolution of the autoreactive phenotype is impeded in the absence of T_FR_. (A) Overview of experimental timeline: T_FR_-suLicient mice were treated with R848 for 4 weeks, followed by treatment cessation for another 5-6 weeks (10-11 weeks total). Subsets of mice were analyzed at various time points throughout the experiment. (B) Spleen weight to bodyweight ratio of T_FR_-suLicient mice before (week 0), during (weeks 2, 3) or after cessation of R848 treatment (weeks 5, 6, 7, 11). Individual datapoints and mean ± SD are indicated. P-values were computed on log-transformed data using one-way ANOVA with Dunnett’s multiple comparisons test. (C) As (B) but for autoantibody measurements (anti-dsDNA IgG2c). P-values were computed using Kruskal-Wallis test with Dunn’s multiple comparisons test. (D) Overview of experimental timeline: as A but including both T_FR_-suLicient and T_FR_-deficient mice. Analysis after 4 weeks of R848 treatment (week 5), and 6-7 weeks after treatment cessation, compared to untreated controls of each genotype. (E) Spleen weights. Individual datapoints and mean ± SD indicated for n = 6-12 mice per group, pooled from 5 cohorts. To meet normality requirements for statistical analysis, data were log-transformed before p-values were computed using two-way ANOVA with uncorrected Fisher’s LSD multiple comparisons test. (F) Anti-dsDNA (left) and total (right) IgG2c levels. Individual datapoints and mean ± SD for n = 4-12 mice per group, pooled from 5 cohorts. To meet normality requirements for statistical analysis, p-values were computed using two-way ANOVA on log-transformed data (anti-dsDNA IgG2c) or raw data (total IgG2c) with uncorrected Fisher’s LSD multiple comparisons test. (G and H) GC B cell, and PB and PC frequencies determined by flow cytometric analysis of the spleen and inguinal lymph nodes (IngLNs). Individual datapoints and mean ± SD shown for n = 5-12 mice per group, pooled from 5 cohorts. To meet normality requirements for statistical analysis, p-values were computed using two-way ANOVA on log-transformed data with uncorrected Fisher’s LSD multiple comparisons test. (I-L) Outlier analyses of spleen weights, serum anti-dsDNA IgG2c, and GC B cells, PBs and PCs in the spleen. Violin plots show the distributions, and outliers identified using 10% ROUT method are marked in red, representing mice that exhibited a sustained response. The stacked bar graph summarizes the frequency of identified outliers (red, sustained immune response) versus non-outliers (light grey, resolved immune response). (M) Pie charts representing total outlier (= mice with sustained response, despite cessation of inflammatory drive) frequencies summarized over all experimental readouts in Fig 3I-L, for T_FR_-suLicient and T_FR_-deficient mice, respectively. Mice marked as outliers in more than one readout by 10% ROUT analysis were counted only once.

We next assessed the impact of T_FR_ deficiency on resolution of the autoreactive response. T_FR_-sufficient and deficient mice were treated with R848 for 4 weeks and either analyzed immediately in week 5 or rested for 6-7 weeks to be analyzed in week 10/11 (Fig. 3D). Compared to untreated controls, both T_FR_-sufficient and deficient mice developed significant splenomegaly after 4 weeks of R848 treatment (Fig. 3E). However, all T_FR_-sufficient mice displayed reversal of the splenomegaly 6-7 weeks after R848 treatment cessation, whereas T_FR_-deficient mice, to the contrary, did not fully revert their splenomegaly (Fig. 3E). Anti-dsDNA IgG2c levels in sera behaved similarly (Fig. 3F, left panel), and this effect was specific to autoantibodies, because by contrast, total IgG2c levels reverted equally in both groups (Fig. 3F, right panel). Whereas GC B cell levels largely reverted, an impaired phenotype reversal in T_FR_-deficient mice was also evident for PB and PC in both spleen and IngLN (Fig. 3G and H, gating strategy in Fig. S1E). Taken together, these data suggested that reversal of autoreactivity following inflammation resolution was impaired in the absence of T_FR_.

### A substantial fraction of T_FR_-deficient mice shows a sustained autoreactive response

To further investigate whether inflammation triggered a sustained immune response in the absence of T_FR_, we applied a ROUT outlier analysis to identify mice that showed a sustained immune response after the rest period. In the T_FR_-deficient group, several mice showed markedly elevated spleen weight (Fig. 3I) and autoantibody levels (Fig. 3J). GC B cell readouts did not present any outliers (Fig. 3K), whereas PBs and PCs displayed another outlier in the T_FR_-deficient group (Fig 3L). We next looked across all readouts. To avoid inflating outlier frequency, each mouse flagged by the 10% ROUT method was counted only once, even if it appeared in multiple readouts. Yet almost half (43%) of the T_FR_-deficient mice exhibited signs of a sustained immune response after cessation of TLR7 stimulation, whereas all T_FR_-sufficient mice appeared to have reestablished homeostasis (Fig. 3M). This difference in proportions (0/5 and 3/4 affected/unaffected for T_FR_-sufficiency and T_FR_-deficiency, respectively) was significant by one-sided Chi-square test (p=0.0455). Moreover, a binomial test of the observed T_FR_ data (3/4) against the expected distribution (0/7) under the assumption that all mice would revert (as observed in the T_FR_-sufficient group) was highly significant (p<0.0001).

An aggregated outlier analysis across all datasets presented in Figures 1 and 3 additionally identified more outliers in T_FR_-deficient than in T_FR_-sufficient mice based on 10% ROUT analysis of spleen weight, autoantibody levels, total IgG2c levels, and GC B, PB and PC frequencies in the spleen (Fig. S3A-E). The summary analysis of all immunological measurements across all cohorts revealed that marked deviations in the inflammatory response primarily occurred in T_FR_-deficient mice (19% out of 36 mice vs. 3% out of 29 mice in the control group, p=0.0255 by one-sided Chi-square test) (Fig. S3F and G).

Taken together, these findings suggested a failure to maintain and reestablish immune homeostasis in a significant fraction of T_FR_-deficient mice during and following cessation of TLR7-driven inflammation.

**Figure S3.**
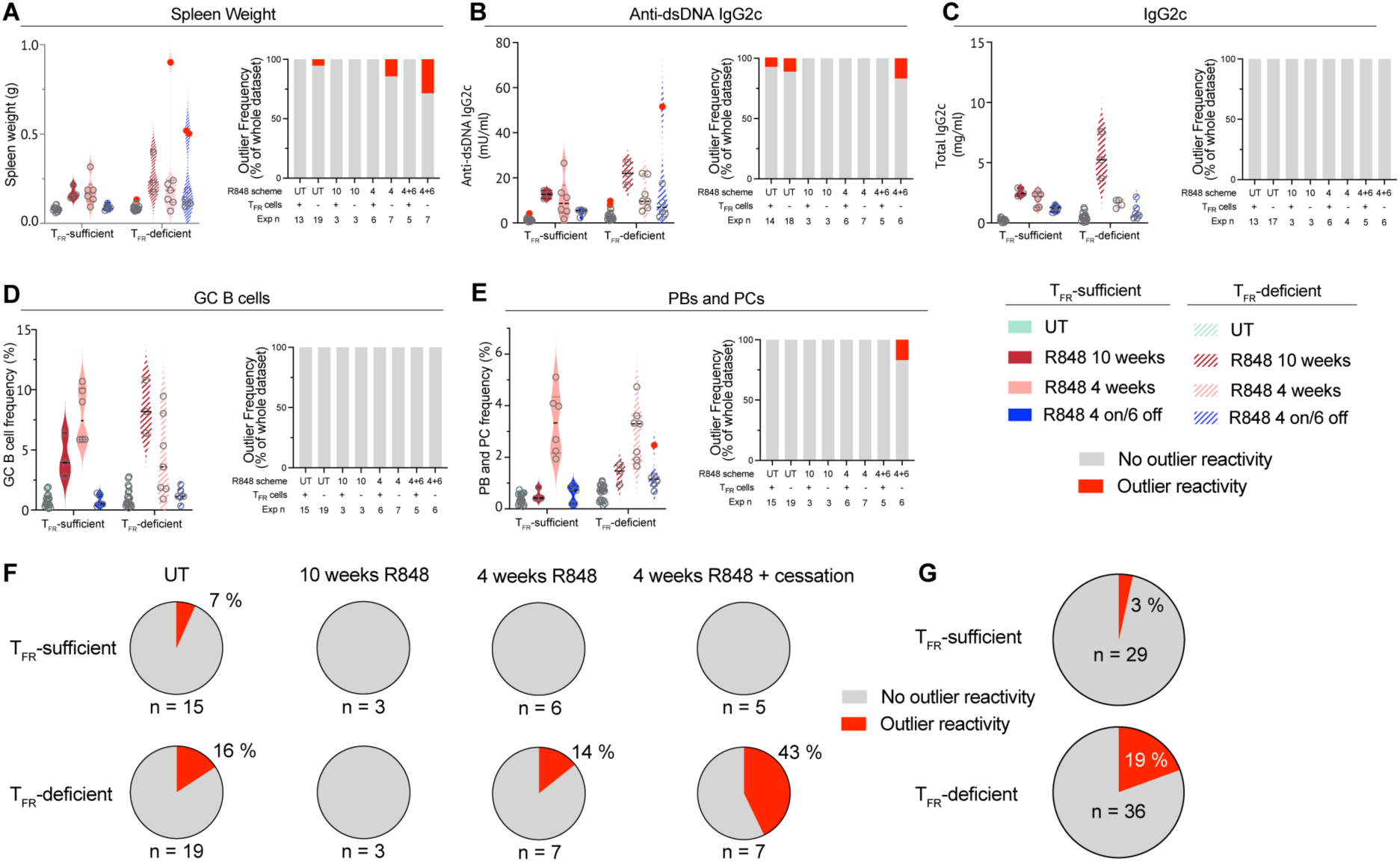
Detailed outlier analysis highlights an enhanced and sustained autoreactive response in T_FR_-deficient mice. (A-E) Outlier analyses of spleen weights, anti-dsDNA IgG2c, total IgG2c, GC B cells, PBs and PCs in the spleen. Violin plots show the distributions, and outliers identified using 10% ROUT method are marked in red, representing mice that exhibited an enhanced or sustained response. The stacked bar graphs summarize the frequency of identified outliers (red, enhanced/sustained immune response) vs. non-outliers (light grey, normal/resolved immune response). (F) Pie charts representing outlier frequencies for T_FR_-suLicient (top) and deficient (bottom) groups, based on readouts in Fig. S3A-E within each of the four experimental conditions (UT, 10 weeks R848, 4 weeks R848, 4 weeks R848 treatment followed by 6 weeks rest) represented in main Figures 1A, 3A, and 3D. Each pie chart shows the relative proportion of mice with a heightened immune response (red segment, outlier response) and non-outliers (light grey segment). Mice that were marked as outliers in 10% ROUT outlier analysis were counted only once, even if they were outliers in several readouts. (G) As F, but all mice were pooled within T_FR_-suLicient and deficient groups.

### T_FR_ respond dynamically to limit spontaneous GC expansion, not GC contraction

These results indicated a role for T_FR_ in controlling the response to inflammation, but since both T_REG_– and T_FH_-derived T_FR_ depend on Bcl6 and express FoxP3, both populations would be affected by our genetic T_FR_ blockade strategy. However, T_FR_ derived from thymic T_REG_ precursors would be expected to emerge early during the response, whereas T_FH_-derived T_FR_ would be predicted to arise later through phenoconversion of T_FH_ in end-stage GCs. To distinguish the contribution of these populations to control and reversibility of inflammation-driven GC responses, we next examined the temporal window during which follicular regulatory activity operates. To this end, we employed an abdominal imaging window (AIW) ^44^ and serial intravital two-photon microscopy to track T_FR_ dynamics within individual splenic GCs during initiation and resolution of the TLR7-driven response (Fig. 4A-C). GCs were visualized by intravital labelling of FDC networks and we leveraged a FoxP3-GFP-DTR line as a FoxP3 reporter. GCs were identified as FDC networks in conjunction with highly vacuolar tingible body macrophages (TBMs). 4D reconstruction of the FDC network (Fig. S4A-C) revealed that GCs were initiated and expanded during R848 treatment (Fig. 4D and E), whereas they contracted following cessation of R848 treatment (Fig. 4F and G). Quantitative 4D image analysis confirmed that FDC networks expanded progressively to almost double their baseline size over 13 days of R848 treatment (Fig. 4H, Fig. S4D), whereas they contracted to almost half their peak size over 13 days following treatment cessation, returning them to their original baseline (Fig. 4I, Fig. S4D). Regression analyses revealed an average increase in FDC network size of around 8.3% per day during R848 treatment, whereas the average contraction rate following R848 cessation was 3.4% per day. The slopes differed significantly (p = 0.0009) (Fig. S4E). Despite substantial inter-GC variability, all GCs expanded during R848 treatment and contracted following treatment cessation (Fig. S4F). The heterogeneity between individual GCs was more evident during the initiation of the response (Fig. S4E and F).

**Figure 4.**
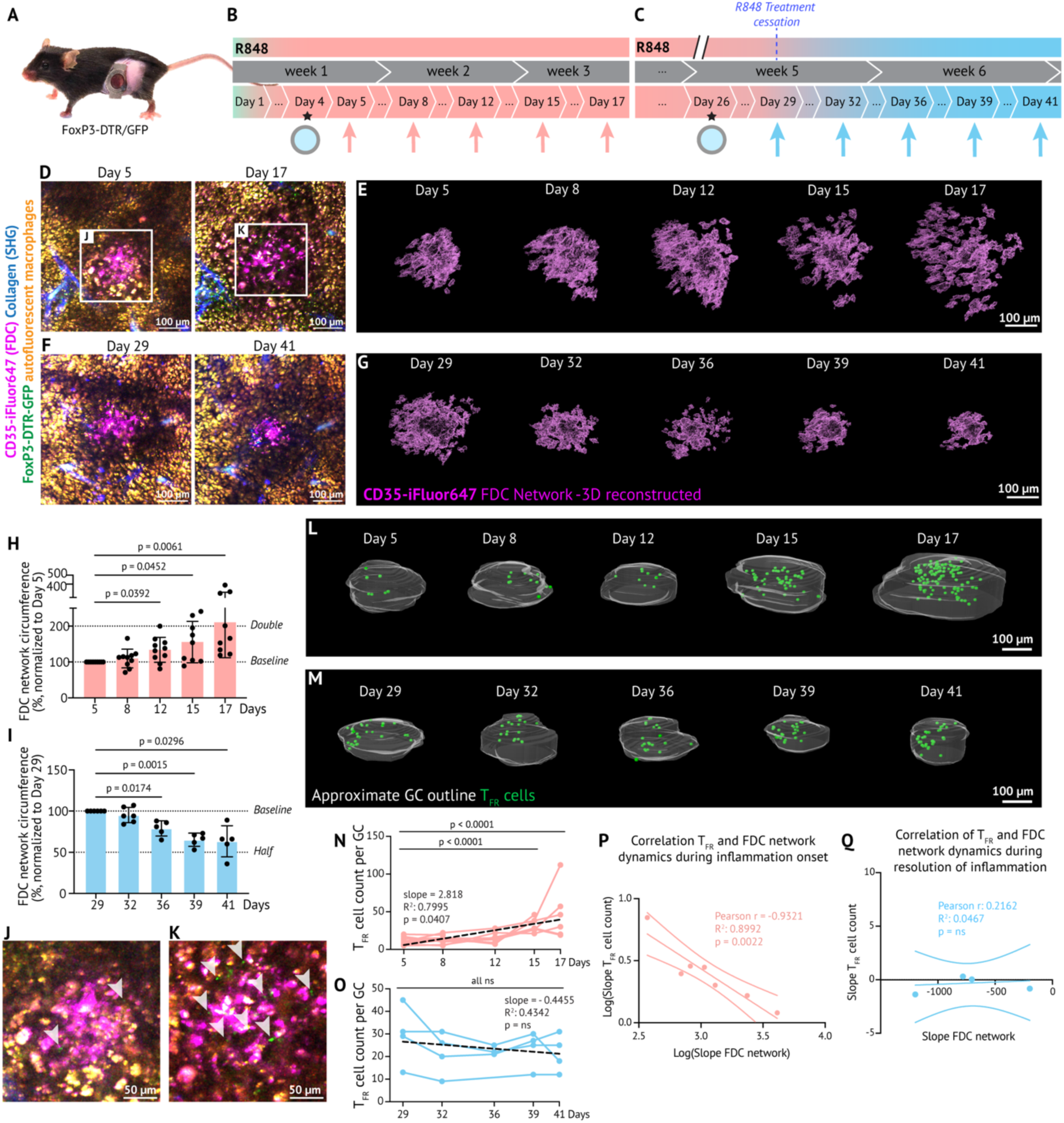
T_FR_ regulate the expansion, not contraction, of individual GCs in response to inflammation. (A) Representative image of a mouse (FoxP3-DTR/GFP) with an AIW implant for serial intravital imaging of the spleen. (B and C) Overview of experimental setup. Day 1 marks the first R848 treatment; day 29 marks the first day after R848 treatment cessation. The window and asterisk mark the surgical implantation of the AIW, and arrows mark imaging days (5x imaging over two weeks per mouse). (D) Micrographs of a GC during R848 treatment, from the first imaging day (day 5, left) and last imaging day (day 17, right). Micrographs show comparable 2D imaging planes at around 110 µm below the spleen capsule and have been median filtered, background subtracted, 3D maxima filtered, and linearly adjusted in brightness and contrast. Scale bar: 100 µm calculated by pixels, as precise scaling of image is not possible after 3D maxima filtering). (E) Representative 3D-reconstructed FDC network (top view) from each imaging day during R848 treatment. Scale bar: 100 µm. (F) As D, but visualizing a GC after cessation of R848 treatment, from the first imaging day (day 29, left) and last imaging day (day 41, right). (G) Representative 3D-reconstructed FDC network (top view) from each imaging day after R848 treatment cessation. Scale bar: 100 µm. (H) FDC network size of individual GCs over the course of R848 treatment. Analysis was done on 2D average Z-projections of an entire volumetric image stack and measurements were normalized to day 5 (= 100% baseline size). The data represent means ± SD from 10 GCs, pooled from 3 mice (2-4 GCs per mouse) across 2 independent cohorts. One GC was excluded on day 15 due to loss of the field of view caused by extensive tissue remodeling. P-values were computed using one-way ANOVA with a mixed-eLects model and Dunnett’s post hoc test, comparing all imaging days to the baseline day 5, using log-transformed data to meet normality requirements for statistical analysis. (I) FDC network size of individual GCs following cessation of R848 treatment, normalized to day 29 (= 100% peak size). The data represent means ± SD from 6 GCs, pooled from 3 mice (2 GCs per mouse) across 2 independent cohorts. P-values were computed using one-way ANOVA with a mixed-eLects model and Dunnett’s post hoc test, comparing all imaging days to day 29. (J and K) Magnified views from micrographs in D showing T_FR_ (indicated by white arrows) in GCs on days 5 and 17 of R848 treatment, respectively. Scale bar: 50 µm, calculated by pixels, as precise scaling of image is not possible after 3D maxima filtering. (L and M) 3D rendering of an entire GC (side view) over two weeks during R848 treatment and after treatment cessation, respectively. T_FR_ are rendered as green spheres (visualized larger than actual cell bodies). Scale bar: 100 µm. (N and O) T_FR_ cell numbers in individual GCs during R848 treatment and after treatment cessation, respectively. In N, n = 9 GCs from 3 mice in 2 cohorts. Two were excluded from day 17, 1 from day 15, and 1 from day 12 due to compromised imaging conditions or spatial overlap of PT tracks (T_REG_) and GC (T_FR_). In O, n = 4 GCs from 3 mice in 2 cohorts. P-values were computed by two-way ANOVA main eLects analysis and Fisher’s LSD post-hoc test, comparing all imaging days to the baseline day 5 on log-transformed and day 29 on non-transformed data, respectively. (P and Q) Change in FDC network size vs. change in T_FR_ numbers for individual GCs during R848 treatment and after treatment cessation, respectively (data in P were log-transformed). Each point represents longitudinal data from one GC, with n = 7 GCs from 3 mice and n = 4 GCs from 3 mice, respectively. Lines and 95% CI based on linear regression analysis (see also Figure S5E and F for more information).

**Figure S4.**
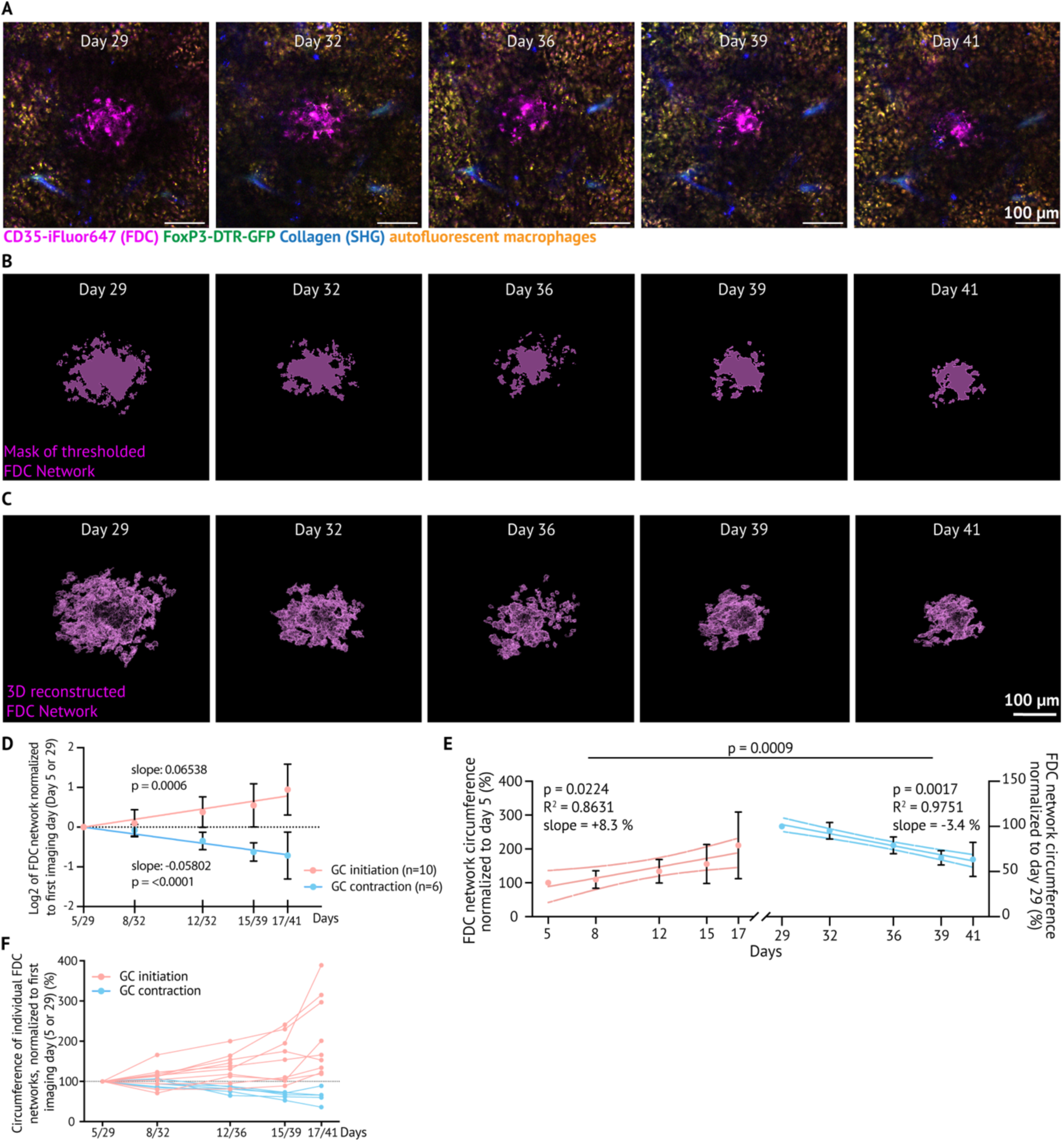
Representative workflow and additional 4D imaging analyses of FDC network dynamics. (A) Serial intravital imaging of a GC over 2 weeks after R848 treatment cessation. The micrographs were standardized in planes to show comparable 2D imaging frames at around 110 µm ± 10 µm below spleen capsule. For representative purposes micrographs were median filtered and adjusted in contrast (linear). Scale bar: 100 µm. (B and C) Image processing pipeline to 3D-reconstruct the FDC network. After image preprocessing in Fiji, a threshold was set over the FDC network using the ‘Big structure detection’ mode in 3D in Arivis. Single image planes showing the thresholded FDC network area are shown in B. Each plane was manually reviewed to ensure inclusion of only the labeled FDC network while excluding autofluorescent macrophages. Thresholds were adjusted when necessary, and the resulting masks were re-evaluated accordingly. In C, the final 3D-reconstructed FDC network is shown in top view. Scale bar: 100 µm. The respective imaging planes from the selected micrographs in A correlate with the selected thresholded masks in B, and 3D reconstruction in C. (D) FDC dynamics during R848 treatment and after treatment cessation. FDC network circumference (measured on the 2D Z-projection) was normalized to the first imaging time point (day 5 and 27, respectively) and log2-transformed to visualize fold change in size over time. Slopes were calculated using simple linear regression based on daily means. Displayed are means ± SD, based on n = 6 or 10 GCs from 3 mice per group, pooled from 2-3 independent cohorts. Two imaging days of one GC during R848 treatment were excluded due to compromised imaging conditions. (E) Linear regression analysis of FDC network circumference changes during R848 treatment and after cessation, normalized to day 5 or 29, respectively. The 95% CIs for the regression lines are indicated by the dashed lines. (F) Individual FDC network dynamics during R848 treatment or after treatment cessation. Same imaging datasets as in D, but in this graph, individual GCs are displayed, revealing individual diLerences in the slopes of FDC network remodeling, especially in the GC initiation group.

The greater variability observed during GC initiation, compared with the more tightly distributed contraction phase, suggested the potential presence of a regulatory bottleneck early in the response. We therefore examined the temporal dynamics of T_FR_ to assess whether their activity constitutes a rate-limiting step in the regulation of incipient GCs (i.e., T_REG_-derived T_FR_) or end-stage GCs (i.e., T_FH_-derived T_FR_) in response to inflammation. T_FR_ were identified as GFP^+^ cells within the follicle (Fig. 4J and K) and 3D reconstruction of entire GCs visualized the fluxing T_FR_ numbers associated with inflammation-driven GC expansion and contraction (Fig. 4L and M, respectively). Image analysis revealed that T_FR_ numbers consistently increased during the R848 treatment as GCs expanded (Fig. 4L and N), in agreement with our previous observation (Fig. 1L). Conversely, T_FR_ numbers stagnated and declined after cessation of the inflammatory stimulus, as GCs contracted (Fig. 4M and O). To corroborate our imaging-based readouts of T_FR_ dynamics, we performed flow cytometric analysis of T_FR_ frequencies in spleens before (day 0), immediately after 4 weeks of R848 treatment (day 29), and 2 weeks after stopping treatment (day 44) (gating strategy in Fig. S5A). This revealed a robust, around five-fold, and statistically significant increase in T_FR_ following 4 weeks of treatment, followed by a decrease after the 2-week rest period (Fig. S5B). T_REG_ frequencies also changed significantly, albeit with only an approximately two-fold difference at day 29 (Fig. S5C). Unlike T_REG_, T_FR_ are known to downregulate the alpha-chain of the high-affinity IL-2 receptor, CD25 ^41,45,46^. In support of their T_FR_ identity, gated T_FR_ showed markedly lower CD25 expression than T_REG_ at the height of the GC response at day 29. T_FH_ did not express CD25 on day 29 (Fig. S5A and D).

**Figure S5.**
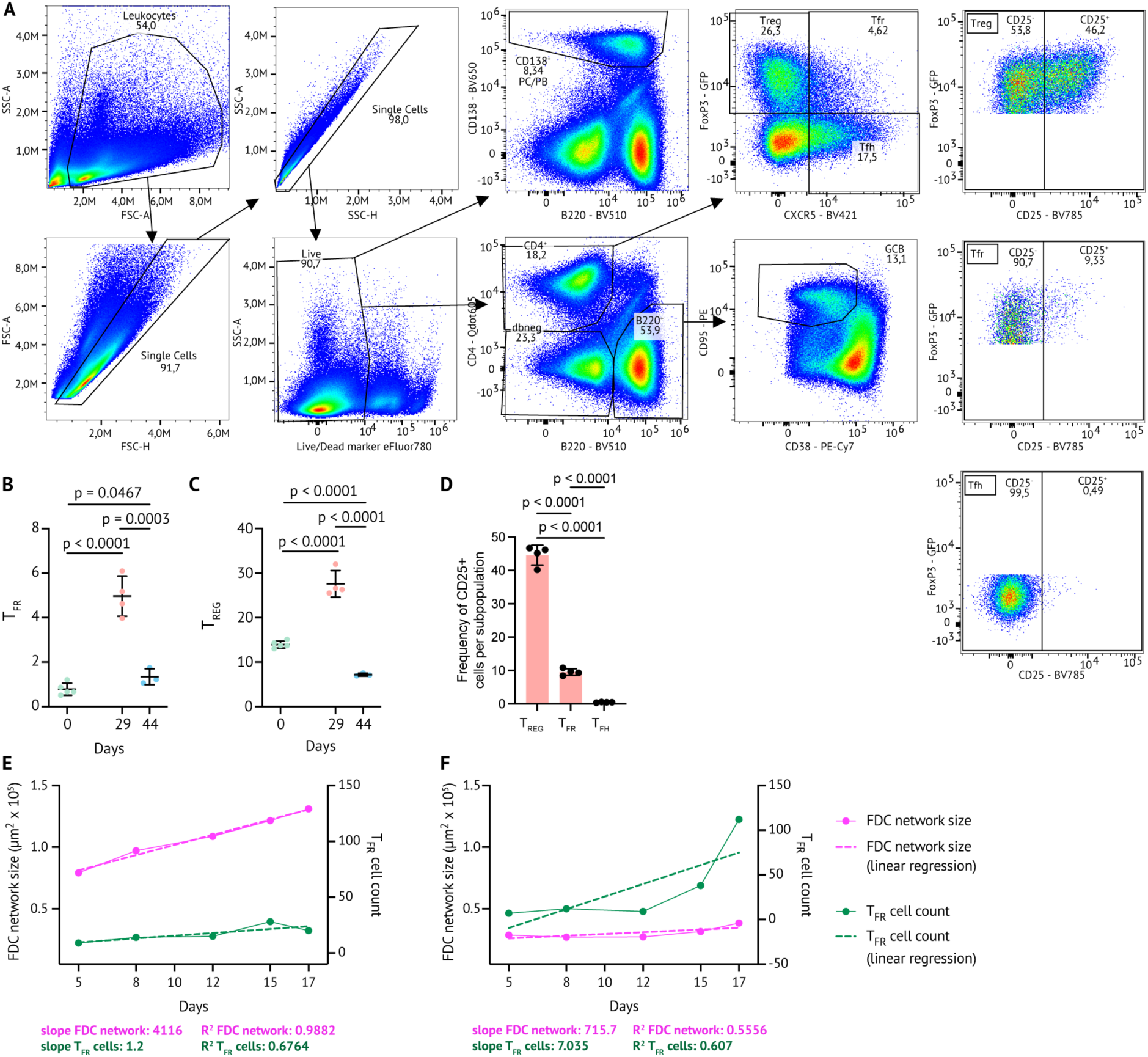
Additional information on T_FR_ cell dynamics. (A) Gating strategy. (B and C) Respectively T_FR_ and T_REG_ frequencies in spleens of untreated FoxP3-DTR-GFP mice (Day 0; n = 5), after 4 weeks of R848 treatment (Day 29, n = 4), and 2.5 weeks after R848 treatment cessation (Day 44, n = 3). P-values were computed using one-way ANOVA with Tukey’s multiple comparisons test on log-transformed data to meet normality requirements for statistical analysis. (D) Frequency of CD25^+^ cells amongst T_REG_, T_FR_ and T_FH_. P-values were computed using one-way ANOVA with Tukey’s multiple comparisons test with pooled variance on log-transformed data to meet normality requirements for statistical analysis. (E) The changes in FDC network size, measured on a 2D Z-projection (magenta, left y-axis), and changes in T_FR_ counts (green, right y-axis) over time within a single GC, with linear regression lines (dashed, respective colors). Slopes and R^2^ values are indicated below the graph. This is an example of an FDC network that expanded markedly (high slope) and displayed a low T_FR_ influx rate (low slope). In Figure 4P and Q, the slope values are plotted against each other to assess their correlation. (F) As E, but an example of an FDC network that did not expand very much (low slope) and displayed a high T_FR_ influx rate (high slope).

Based on our imaging data, we quantified the rates of change in FDC network size and T_FR_ numbers for each GC (representative examples in Fig. S5E and F). We then assessed the relationship between these parameters by plotting the slope of FDC network size versus the slope of T_FR_ counts for each GC during R848 treatment (Fig. 4P) and after R848 treatment cessation (Fig. 4Q). This uncovered an inverse correlation of the change in T_FR_ and the expansion of FDC network, demonstrating that the higher the increase in T_FR_ during the inflammatory stimulus, the less the GC expanded, and vice versa (Fig. 4P). In contrast, T_FR_ stagnated or declined in concert with the GC contraction after the cessation of the inflammatory stimulus (Fig. 4Q). Overall, these results suggested that T_FR_ limit the establishment and early expansion of spontaneous GCs in response to inflammation, whereas they did not expand during the GC contraction, as would have been expected if T_FH_ phenoconverted to drive this process.

### T_FR_ derive from extrafollicular precursors during spontaneous GC establishment, and most express Helios

As T_FH_ are not yet abundant during GC initiation, early T_FR_ most likely originate from T_REG_ precursors. Hence, the early increase in T_FR_ and the inverse correlation of the number of T_FR_ with the magnitude of GC expansion (Fig. 4N and P), suggested that the T_FR_ controlling spontaneous GC responses to inflammation were bona fide thymic T_REG_-derived T_FR_. We noted an increase in GFP^+^ cells on perivascular T-tracks (PT-tracks) (Fig. 4D, GFP^+^ cells on blue SHG-signal in lower left corner) previously reported to serve as unidirectional migration paths for T cells from the red pulp via bridging channels towards T zones ^47^. This suggested T_REG_ might be migrating into the follicle and differentiating into T_FR_ in response to the inflammatory stimulus. To investigate this, we sorted out CD4^+^ T cells from FoxP3-DTR/GFP donors by magnetic-activated cell sorting, then FACS sorted FoxP3-GFP^+^ (T_REG_) and FoxP3-GFP^−^ (non-T_REG_) cells. The non-T_REG_ cells were labelled with Qtracker Qdot655 for long-term tracking. GFP^+^ T_REG_ and Qdot655^+^ non-T_REG_ cells (Fig. S6A and B) were co-transferred into FoxP3-Cre;Bcl6^flx/flx^ recipients, which had begun R848 treatment two days prior, and treatment was continued for another two weeks (Fig. 5A). The day before and on the day of harvest, mice were labelled intravitally with anti-CD35-A647 and anti-CD169-PE, respectively, then spleens were harvested for analysis by flow cytometry and two-photon explant imaging. By flow cytometry (gating strategy in Fig. S6C), we observed a low frequency of T_FR_, the majority of which (∼75%) were Helios^+^, similar to T_REG_ (Fig. 5B). Fixation and permeabilization for intracellular flow staining destroy endogenous fluorophores, requiring staining with anti-GFP-A488 to identify endogenous fluorophore-positive cells. Because donors are GFP^+^ and recipients are YFP^+^, and anti-GFP recognizes both, we could not distinguish their origin using this parameter. However, since FoxP3-Cre;Bcl6^flx/flx^ recipients cannot generate primary T_FR_ nor phenoconvert T_FH_, the observed T_FR_ must have been donor-derived. Moreover, none of these were Qtracker655^+^, which, together with the dominance of Helios^+^ cells in this subset, indicated that T_FR_ were derived from adoptively transferred T_REG_ precursors that had entered incipient GCs. This was confirmed by explant imaging, in which adoptively transferred FoxP3-GFP^+^ cells (endogenous fluorophore signal) localized to both follicles and PALS (Fig. 5C), and were found to be mainly Helios^+^, whereas we did not detect any signal from Qdot655 (Fig. 5D-I). Overall, the flow cytometric and spatial two-photon analyses indicated that early spontaneous GC reactions were controlled by thymic T_REG_-derived T_FR_ rather than by phenoconverted T_FH_ or peripheral T_REG_.

**Figure 5.**
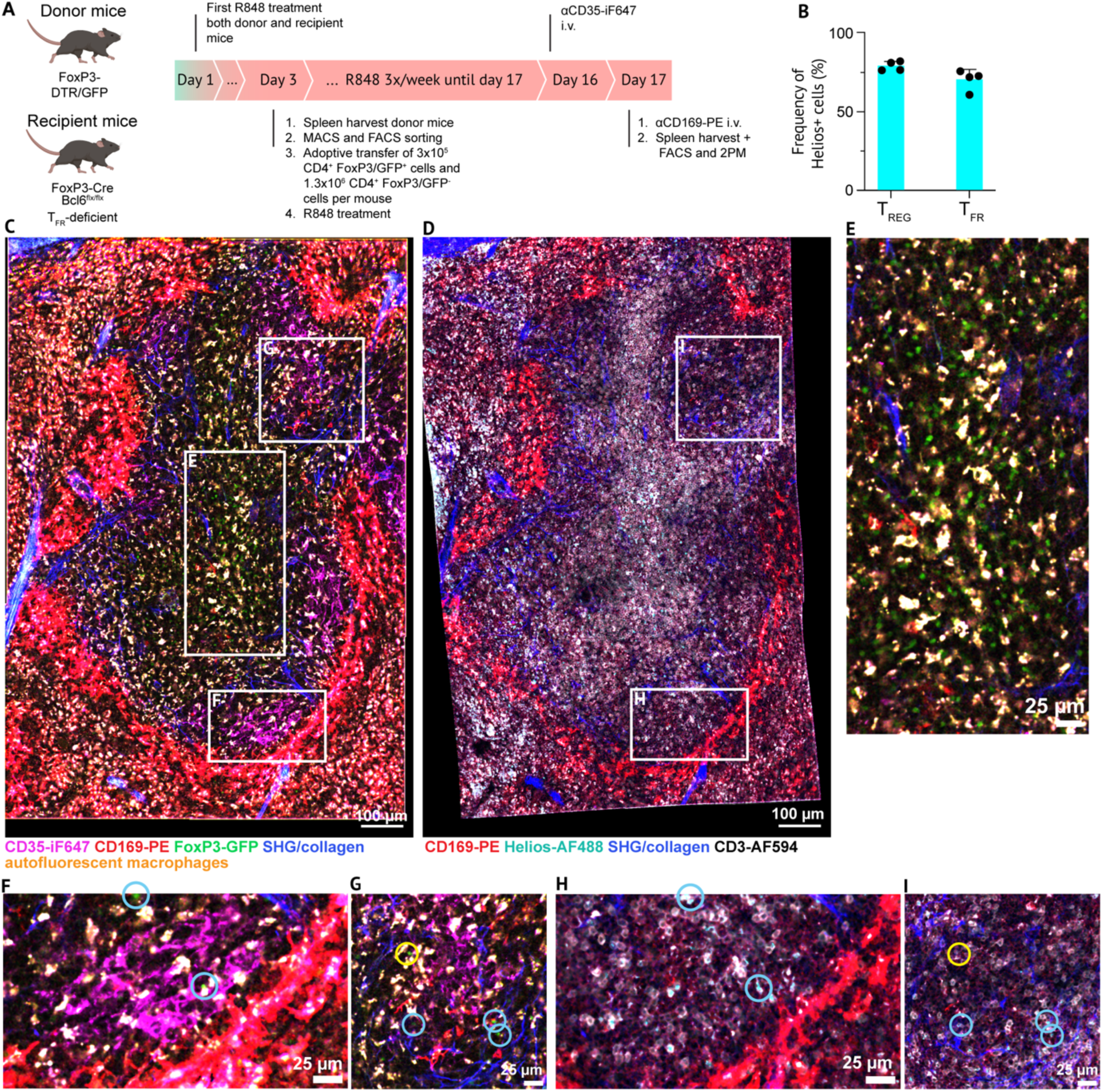
T_FR_ engaging in spontaneous GCs can derive from extrafollicular precursors and most express Helios. (A) Overview of adoptive transfer experiment to investigate T_FR_ provenance. (B) Frequency of Helios^+^ cells among T_REG_ and T_FR_ in T_FR_-deficient mice after adoptive co-transfer of T_REG_ and non-T_REG_ and 2.5 weeks of R848 treatment. Mean of n = 4 mice with 2 technical replicates each. (C) Overview of a white pulp area of a fresh spleen explant from a T_FR_-deficient mouse after adoptive co-transfer of T_REG_ and non-T_REG_ and 2.5 weeks of R848 treatment, showing several FDC networks (CD35-iF647), metallophillic macrophages (CD169-PE), T_REG_ and T_FR_ (FoxP3-DTR/GFP). Scale bar: 100 µm. Representative of n = 4 mice. (D) Same area overview as C, after tissue fixation and staining with anti-Helios-A488 and anti-CD3-AF594 and digital image registration. Scale bar: 100 µm. (E) Magnified view of the T cell zone outlined in C. Color key shown in panel C applies. Scale bar: 25 µm. (F-I) Magnified views of the same two outlined GCs in C (F and G) and D (H and I). Color key shown in panels C and D applies to panel F+G and H+I, respectively. The blue circles indicate T_FR_ (green) in F and G that were identified as Helios^+^ (cyan) in H and I, respectively. The yellow circles in G and I indicate a T_FR_ (green) that was identified as Helios^−^ (no cyan signal). Scale bars: 25 µm. All micrographs in this figure show a single focal plane from a 100 µm 3D z-stack of 2×3 tiles. Stitched micrographs were background subtracted (rolling ball = 50), median filtered (Despeckle), and adjusted for brightness and contrast (linear).

**Figure S6.**
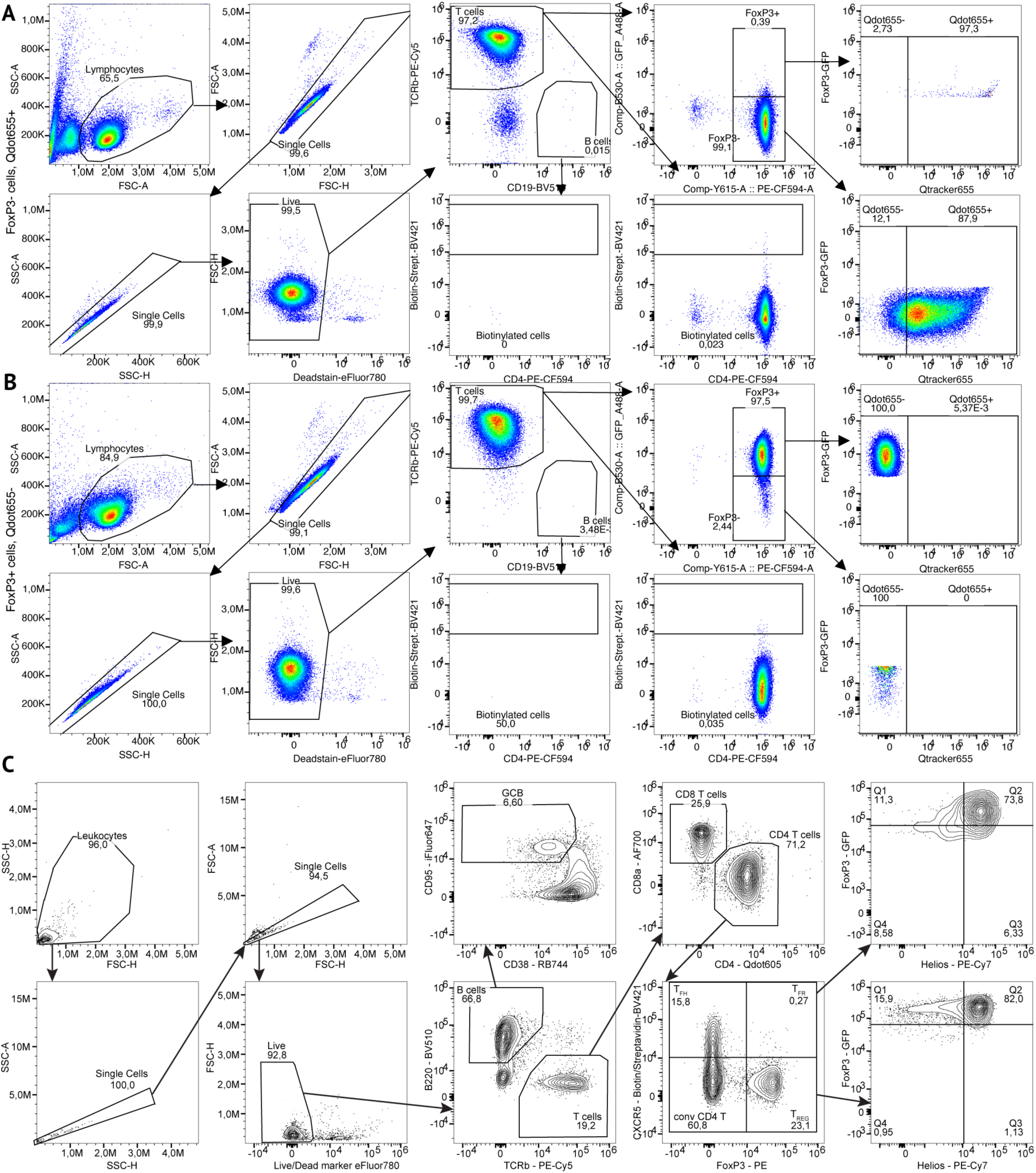
Additional information for adoptive transfer experiment. (A) Flow cytometric analysis of sorted and Qtracker655-labeled FoxP3-GFP– (non-T_REG_) cells immediately before adoptive co-transfer with unlabeled FoxP3-GFP+ (T_REG_) cells. (B) Flow cytometric analysis of sorted FoxP3-GFP^+^ (T_REG_) cells immediately before adoptive co-transfer with Qtracker655-labeled FoxP3-GFP^−^ (non-T_REG_) cells. (C) Gating strategy of the flow experiment analyzing recipient spleens at the end of the experiment.

### T_FR_ controlling spontaneous GCs are transcriptionally similar to and show repertoire overlap with thymic T_REG_

To further evaluate the origin and identity of the T_FR_ that restrain inflammation-driven GC responses, we sorted splenic T_REG_, T_FR_, and T_FH_ from R848-treated FoxP3-GFP/DTR reporter mice (Fig. 6A, gating strategy in Fig. S7A). Sorted cells were subjected to paired 5’-mRNA and V(D)J single-cell sequencing. Unsupervised clustering revealed 8 subclusters (Fig. 6B), identified based on canonical marker expression (Fig. S7B), and confirmed by overlay of various markers (Fig. S7C-K). The proximity of T_FR_ to thymic T_REG_ (tT_REG_) and T_FH_ in the UMAP suggested their transcriptional similarity, whereas they were more distal to peripheral T_REG_ (pT_REG_) (Fig. 6B). Expanded clones mapped mainly to tT_REG_, T_FR_ and T_FH_ clusters, not pT_REG_ (Fig. 6C). Repertoire analysis revealed a high degree of clonal sharing between T_FR_ and tT_REG_ specifically, with a Morisita-index of 0.271, whereas this was only 0.03 and 0.006 for T_FR_ vs. T_FH_ and T_FR_ vs. pT_REG_, respectively (Fig. 6D). This was also evident based on a clonal trajectory map focused on the T_FR_ cluster (Fig. 6E) and a circle plot, which confirmed higher clonal sharing of T_FR_ with tT_REG_ (black arrows in Fig. 6F), than with either T_FH_ or pT_REG_ (Fig. 6F). Overall, these results supported that T_FR_ that restrain spontaneous GCs in response to inflammation originate from tT_REG_. To confirm the identities of the assigned clusters, we plotted key identity markers (Figure 6G). Common to T_FH_ and T_FR_, *Cxcr5*, *Bcl6*, *Tox2*, *Pdcd1*, *Icos*, and *Maf* defined a follicular program, enabling migration into B cell follicles (Cxcr5), establishment and maintenance of GC identity (Bcl6, Tox2), and sustained T-B interactions (Pd-1, Icos, c-Maf). T_FH_ expressed *Cd40lg*, *Il21*, *Ascl2*, and *Nrp1*, reflecting B cell helper function, with Cd40l and Il-21 driving B cell activation and differentiation, while Ascl2 promotes follicular homing and Nrp1 is associated with activated/helper phenotypes. T_FR_ were characterized by *Foxp3*, *Ctla4*, *Ikzf2*, *Tigit*, and *Sh2d1a*, with regulatory lineage being controlled by Foxp3, Ctla4 and Tigit mediating suppression, Ikzf2 (Helios) supporting tT_REG_ identity, and Sh2d1a (SAP) enabling regulatory interactions within the GC. Finally, non-follicular markers included *Prdm1*, *Il2ra*, *Ccr7*, *Sell*, *Klf2*, and *S1pr1*. These genes reflect T-zone residency and counteract follicular programming, with Ccr7, Cd62l (Sell), Klf2, and S1pr1 maintaining recirculation, and Prdm1 (Blimp-1) and Il2ra actively opposing the Bcl6-driven follicular differentiation pathway. Taken together, the marker profile supported the T_FR_ identity.

**Figure 6.**
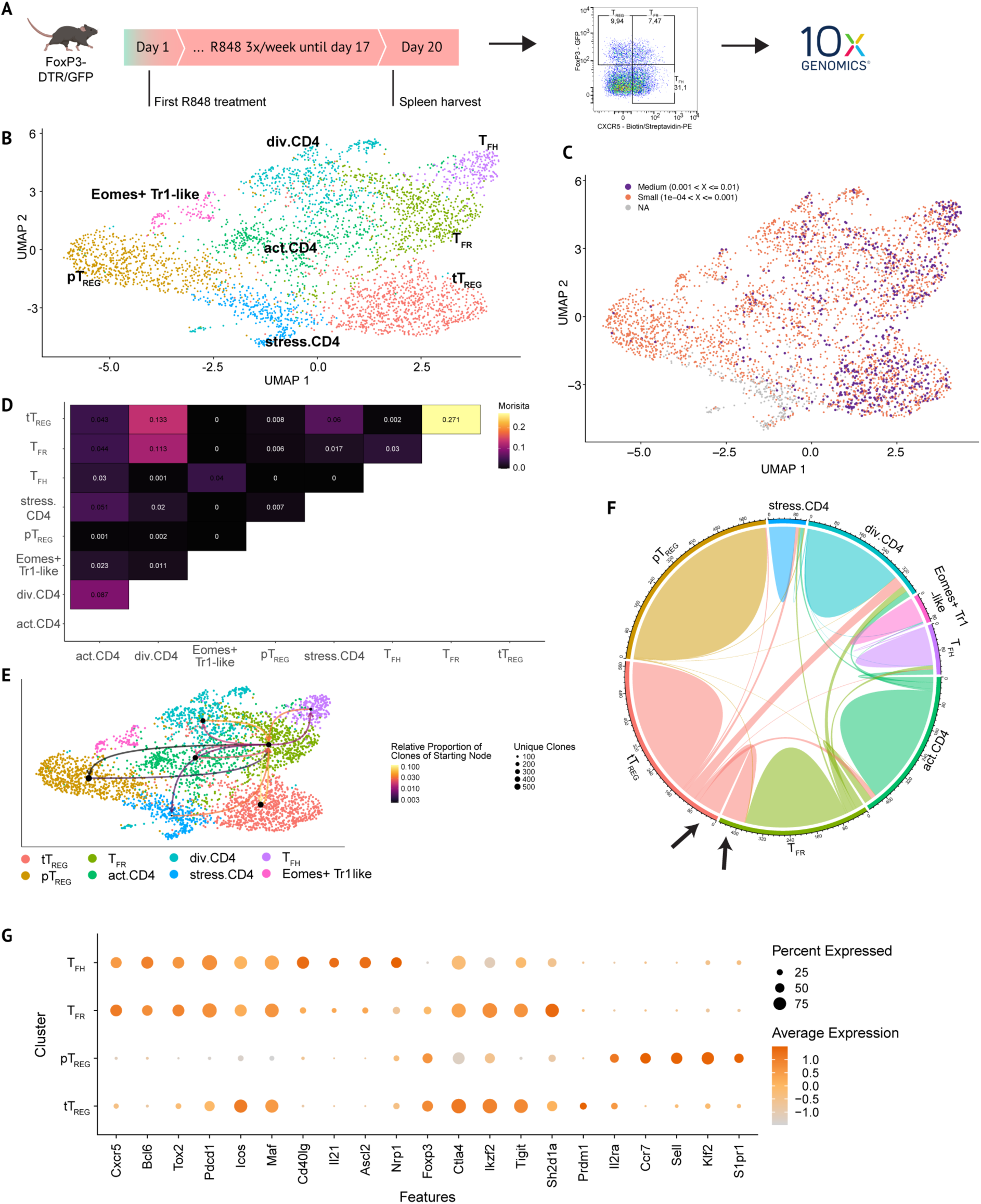
Gene expression profile and TCR repertoire reveal T_FR_ that limit spontaneous GCs originate from tT_REG_. (A) Overview of TCR sequencing experiment to investigate the identity of T_FR_ that restrict early spontaneous GC reactions. FoxP3-DTR/GFP mice were treated with R848 for 2.5 weeks to induce spontaneous GC initiation, then spleens were harvested, and T_FH_, T_FR_, and T_REG_ were FACS sorted and subjected to 10x sequencing. (B) UMAP projection of sorted T_FH_, T_REG_ and T_FR_ (from the spleens of n = 4 mice). Subpopulations are colored by transcriptionally defined clusters. (C) UMAP of sorted T cells, as in B, but colored for clone size. (D) Heatmap showing clonal overlap between transcriptionally defined clusters, quantified using the Morisita index. (E) T_FR_ clonal trajectory overlay on UMAP. (F) Circle plot depicting TCR clonal sharing between populations. Lines connect clusters that share clones, with line thickness proportional to the number of shared clones. Segment size corresponds to the total number of clones in each cluster. Black arrows point to clones shared between tT_REG_ and T_FR_. (G) DotPlot showing expression of key markers discriminating T_FH_, T_FR_, pT_REG_ and tT_REG_.

**Figure S7.**
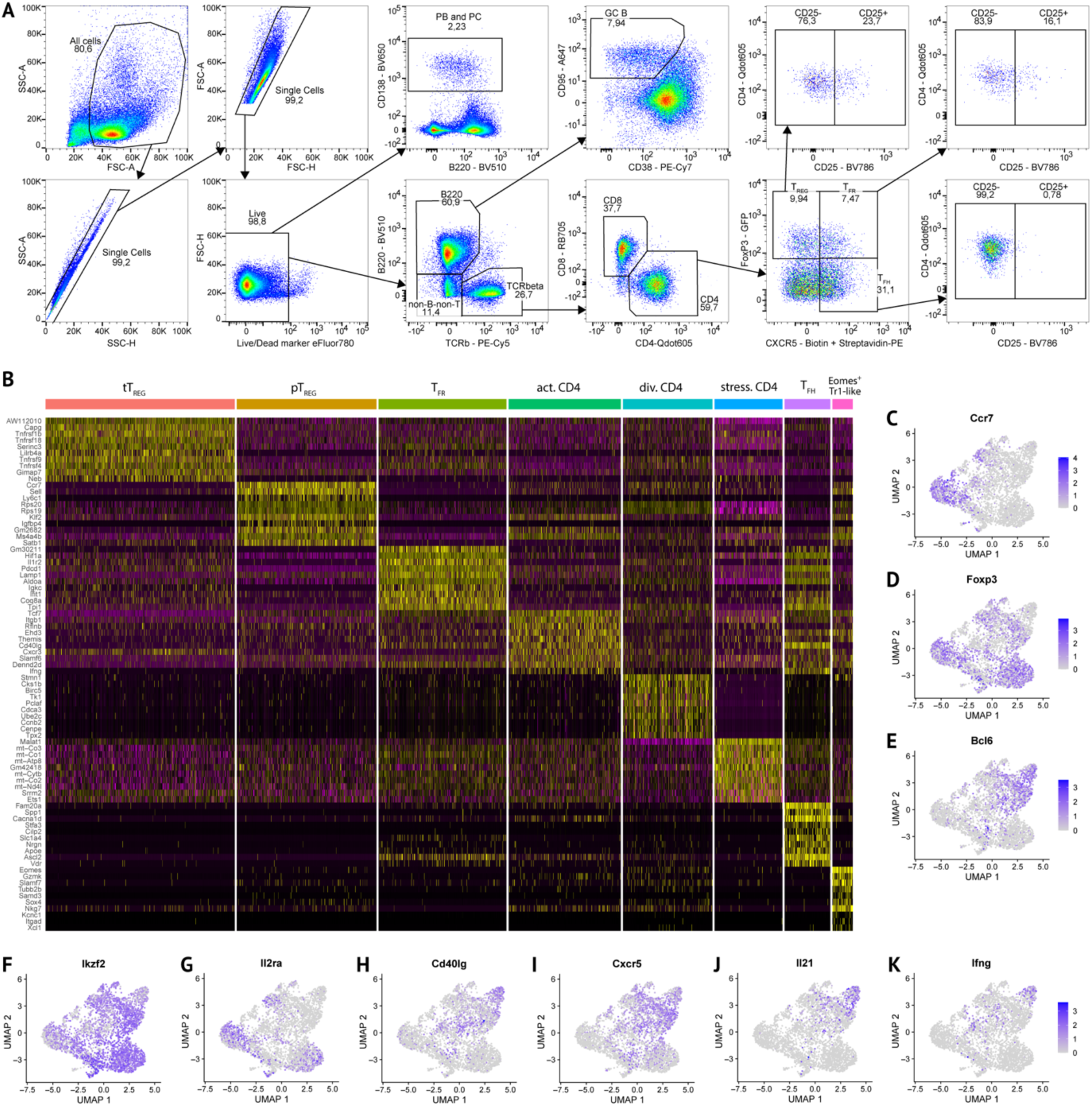
FACS gating strategy and expression markers supporting assigned cell identities. (A) FACS strategy to isolate T_FR_, T_FH_, and T_REG_ for single-cell sequencing. T_FR_, T_FH_, and T_REG_ gates were used for sorting, downstream CD25 gates were only confirmatory of cell identities. (B) Heatmap showing expression of canonical marker genes across clusters. (C-K) Overlay of canonical markers on UMAP from main Fig. 6B.

### Circulating T_FR_ correlate with disease activity scores in SLE patients

T_FR_ that exhibit memory-like features can be found in low numbers in the circulation ^48^. To test whether our observations regarding murine thymic T_FR_ restraining spontaneous GC expansion could be translated to humans, we investigated circulating T_FR_ (cT_FR_) in blood of SLE patients and healthy controls (Fig. 7A, gating strategy in Fig. S8). The cT_FR_ compartment was expanded in SLE patients compared to healthy controls (Fig. 7B), whereas circulating T_REG_ (cT_REG_) and circulating T_FH_ (cT_FH_) were comparable (Fig. 7C and D). We corroborated their respective T_REG_, T_FR_, and T_FH_ identity based on high, intermediate, and low CD25^+^ frequencies, respectively (Fig. 7E). Analysis of PD1 expression indicated an increased activation of all three populations in SLE patients compared to healthy controls (Fig. 7F). The majority (∼80%) of cT_REG_ and cT_FR_ in both healthy controls and SLE patients were Helios positive (Fig. 7G and H). There was a trend towards a higher frequency of Helios positivity among the expanded cT_FR_ in SLE patients, although this did not reach significance (Fig. 7H). We also observed a positive, albeit non-significant, correlation between cT_FR_ levels and SLEDAI score (Fig. 7I, healthy individuals set to –1), indicating that the higher the disease activity, the more the cT_FR_ compartment expanded. The frequency of circulating tT_FR_ (Helios^+^ T_FR_) correlated positively and significantly with the SLEDAI score (Fig. 7J), suggesting that, as we observed in mice, tT_REG_-derived T_FR_ expand during inflammation to constrain spontaneous GC responses in humans.

**Figure 7.**
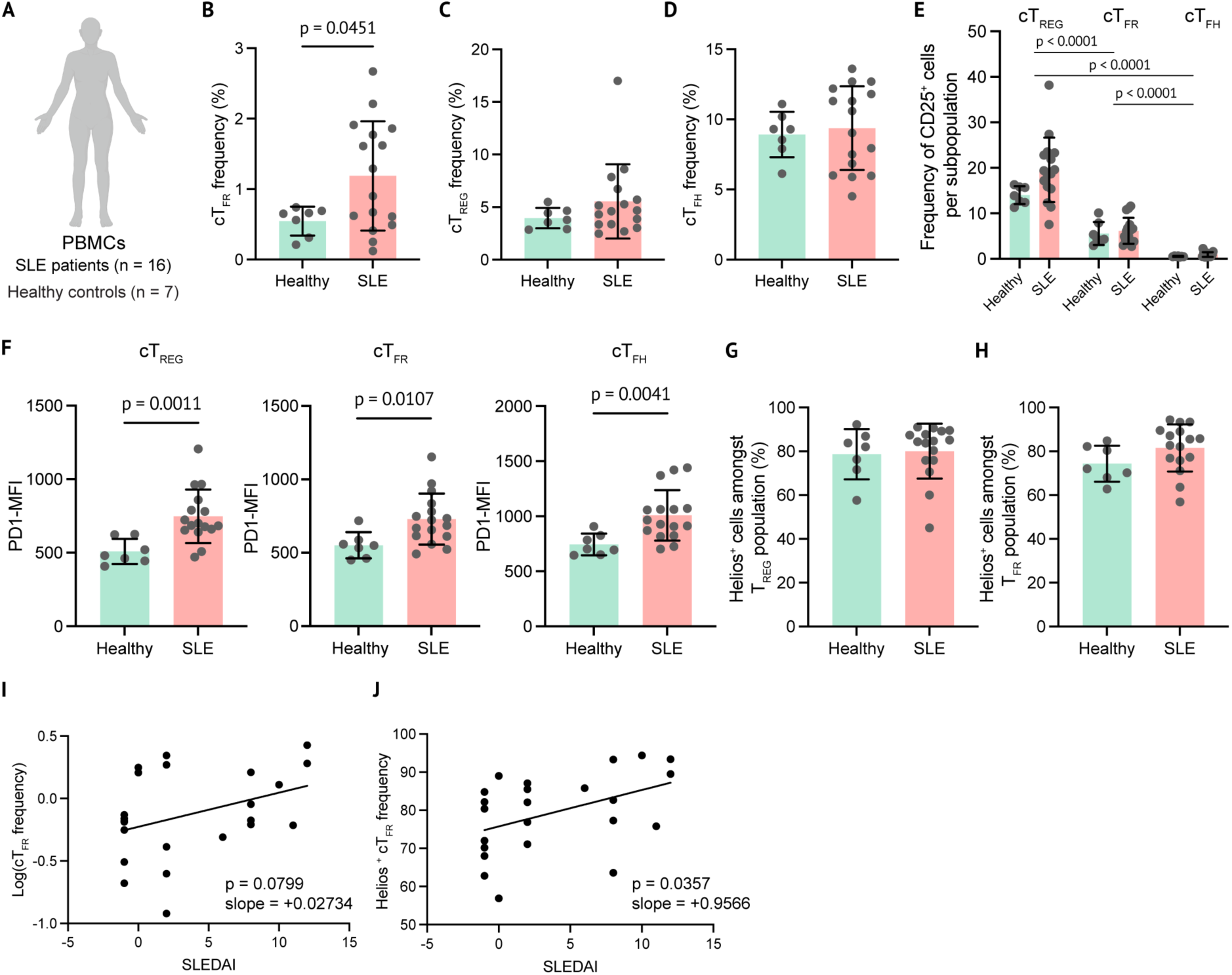
Circulating T_FR_ correlate with disease activity scores in SLE patients. (A) Peripheral blood mononuclear cells (PBMCs) isolated from whole blood samples from 7 healthy controls and 16 SLE patients with varied disease activity scores were subjected to flow cytometric analysis. (B-D) Frequency of cT_FR_, cT_REG_, and cT_FH_, respectively, in healthy controls (n=7) and SLE patients (n=16). Graphs represent cell frequencies out of CD4^+^ T cells, and P-values were calculated using an unpaired t-test. cT_REG_ data were log-transformed to meet normality requirements for parametric test. (E) Frequency of CD25^+^ cells amongst cT_REG_, cT_FR_, and cT_FH_ in healthy controls and SLE patients. Cell frequencies were pooled from SLE patients and healthy controls for each subpopulation, and P-values were calculated using ordinary two-way ANOVA with Tukey’s multiple comparisons test on log-transformed data to meet normality requirements. (F) PD1-MFI of cT_REG_, cT_FR_, and cT_FH_ from healthy controls and SLE patients. P-values were calculated using unpaired t-test on log-transformed data to meet normality requirements for parametric test. (G and H) Helios^+^ cell frequencies amongst cT_REG_ and cT_FR_ in healthy individuals and SLE patients. P-values were calculated using unpaired t-test. (I) Scatterplot of log[cT_FR_ cell frequency] versus SLEDAI score for patients and healthy controls (set to –1). The line represents the linear regression fit, with the slope and p-value for the test of significant deviation from zero. cT_FR_ cell frequencies were log-transformed to meet normality requirements. (J) Scatterplot of circulating thymic T_FR_ cell frequency (Helios^+^ T_FR_ cells) versus SLEDAI score for patients and healthy controls (set to –1). The line represents the linear regression fit, with the slope and p-value for the test of significant deviation from zero.

**Figure S8.**
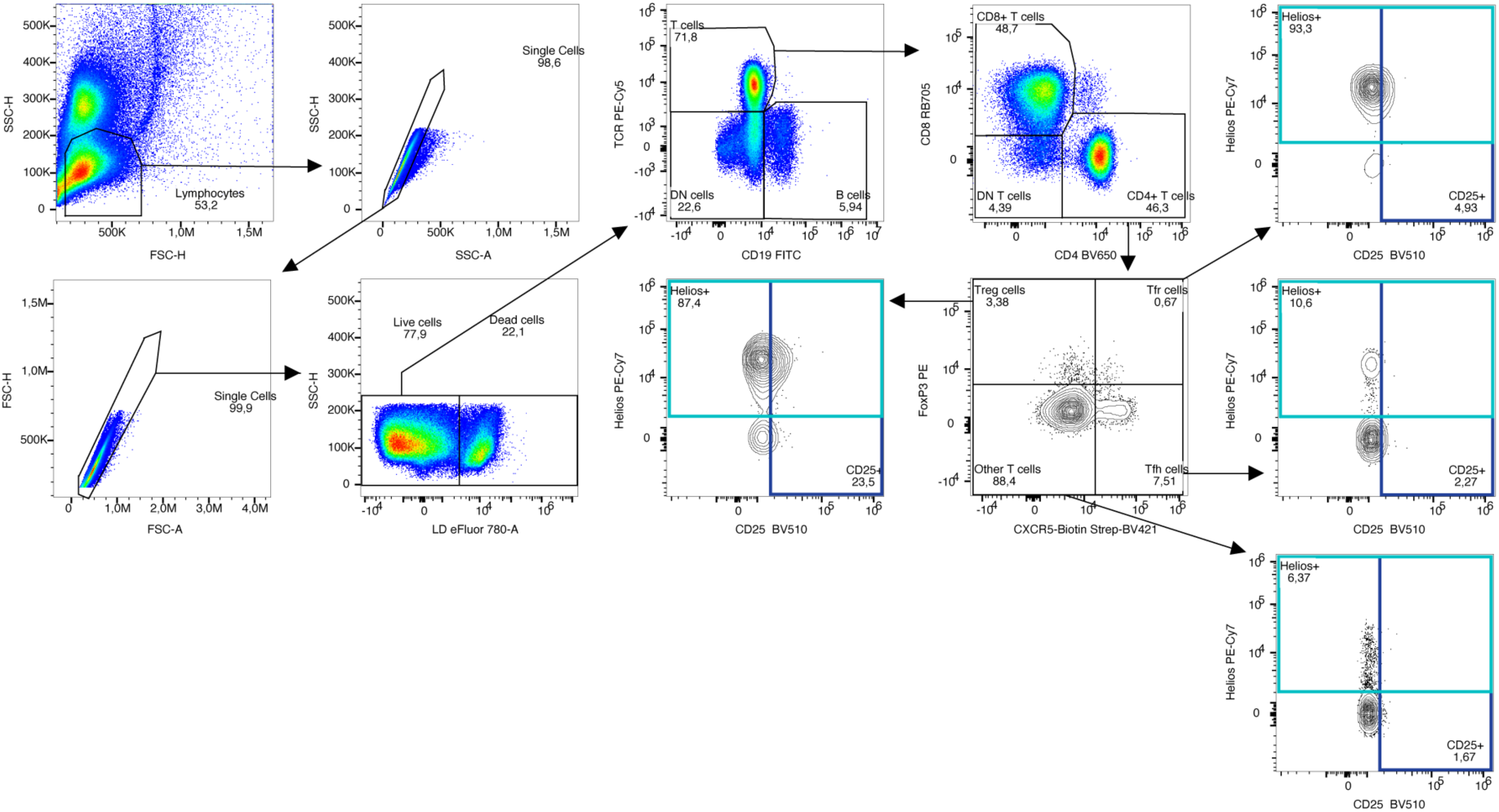
Gating strategy for analysis of PBMCs from healthy individuals and SLE patients. Gating strategy based on T cell panel to identify T_REG_, T_FR_, T_FH_ and define their Helios and CD25 status.

Collectively, our findings indicated that in both mice and humans, tT_REG_-derived T_FR_ play a critical role in protecting against autoimmune disease by limiting the expansion of spontaneous GCs in response to inflammation.

## DISCUSSION

We found that GC regulation by T_FR_ is essential to prevent catastrophic humoral autoimmunity under inflammatory conditions. In the absence of T_FR_, sterile inflammation drove unchecked expansion of autoreactive B and T cell clones, extensive epitope spreading, and irreversible loss of tolerance in a subset of animals. This suggests that GC responses are actively restrained by T_FR_ in response to inflammation. Notably, T_FR_ deficient mice at baseline already displayed some hallmarks of unchecked GC activity but this was exacerbated upon TLR7 stimulation (Figs. 1+2 and S1+S2). This effect was specifically attributable to loss of T_FR_ (Fig. 1M), given that T_REG_ were present in equal or greater numbers in mice with genetic blockade of T_FR_ (Fig. 1L) and FoxP3 TF activity across T_REG_ and other CD4 T cell populations was unaffected by the blockade of T_FR_ (Fig. S2B). Moreover, T_FH_ were present in equal or greater numbers in T_FR_ blocked mice (Fig. 1K), demonstrating that this population was not perturbed by the genetic strategy. Hence, our finding that wild-type mice lost their autoreactive phenotype following R848 treatment cessation, whereas a subset of T_FR_-deficient animals displayed signs of sustained autoreactivity (Figs. 3 and S3) suggests that T_FR_ contribute to maintain tolerance during inflammation. This could play out throughout the response, early during initiation, or when the response should contract again. Our findings from intravital studies (Fig. 4) suggest that the ability to reign in the response once inflammation subsides depends upon early entry of T_FR_ during GC initiation. This aligns with prior findings that T_FR_ can regulate GC B cell development and antigen-specific antibody responses early before GC formation and have less regulatory control after GCs have been initiated ^18^. The early emergence of T_FR_ (Fig. 4) suggested these could not derive from phenoconversion of T_FH_. This was further supported by the ability of FoxP3^+^ T cells sorted from naive SPF donor mice to enter incipient GCs of T_FR_-deficient recipients (Fig. 5), because this adoptively transferred population should be heavily dominated by tT_REG_. Moreover, analyzing T_REG_, T_FH_, and T_FR_ bulk-sorted from R848-treated mice, we observed expanded clones among tT_REG_, T_FH_ and T_FR_, but not pT_REG_ (Fig. 6C), indicating that it was tT_REG_ and their GC counterparts (i.e., tT_FR_) that responded to regulate the inflammatory response. In accordance, we observed significant clonal sharing between tT_REG_ and T_FR_, whereas the overlap with T_FH_ and pT_REG_ was negligible (Fig. 6D-F). Finally, T_FR_ displayed a marker profile commensurate with follicle residency and tT_REG_ origin (Figs. 6G and S7).

Our finding that T_FR_ responding to inflammation are derived from tT_REG_ contrasts with studies showing that T_FR_ can arise from T_FH_ during foreign antigen-driven responses ^21,22^. However, whereas in foreign antigen settings, antigen is transient, by contrast, inflammation-driven autoreactive responses are directed towards ubiquitous self-antigens that cannot be cleared. The model employed by Victora and colleagues ^21^ leveraged a prime-boost strategy in which ovalbumin (Ova) carrier-specific T cells, induced in the presence of adjuvant, later supported NP-hapten-specific adoptively transferred B cells responding to NP-Ova in the absence of adjuvant. This strategy effectively uncouples the GC response from inflammatory signaling, which based on our results would preclude tT_REG_-derived T_FR_ engagement, hence isolating GC control to the T_FH_ population that upregulates FoxP3 in response to limiting antigen availability at the back-end of the GC response. Conversely, our model is based on TLR7-driven inflammation in the absence of foreign antigen, driving early recruitment of tT_REG_-derived T_FR_ into GCs, whereas autoreactive T_FH_ never experience limiting antigen availability, and hence play a minor role in this setting. This notion also reconciles apparently conflicting observations regarding T_FR_-T_FH_ repertoire relationships, whereby T_FR_ have variably been found to display repertoire overlap with either T_REG_ or T_FH_ ^19,20^. Likely, most physiological GC responses would involve both tT_REG_-derived T_FR_, i.e., tT_FR_ (autoantigen specific, responding to initial inflammation) and peripheral T_FH_-derived T_FR_, i.e., pT_FR_ (foreign antigen specific, responding to antigen depletion).

The severity of autoreactive GC responses in the absence of T_FR_ appears to scale with the magnitude of inflammation across physiological settings. Modest spontaneous GCs observed during ageing in clean SPF conditions are associated with limited autoreactivity, whereas higher inflammatory states, such as viral infection, adjuvant-driven autoimmunity, or chronic sterile inflammation, progressively exceed regulatory capacity ^17,41,49,50,51^. Our findings place T_FR_ at a critical intersection between inflammatory tone and humoral tolerance, revealing that tT_REG_-derived T_FR_ are indispensable once inflammatory signals exceed a threshold that permits widespread recruitment of autoreactive clones.

In line with this model, patients with SLE experience episodic flares and remissions ^26^, suggesting that disease activity reflects a dynamic balance between inflammatory drivers and regulatory mechanisms. Circulating T_FR_ have previously been observed in Sjögren’s disease patients and were suggested to correlate with GC activity^52^. In line with this, we observed that circulating Helios⁺ T_FR_ correlated with disease activity in SLE patients (Fig. 7), consistent with a model in which tT_REG_-derived T_FR_ expand in response to inflammatory pressure to restrain autoreactive humoral responses. Together, these findings raise the possibility that selective therapeutic enhancement of tT_REG_-derived T_FR_ could attenuate autoreactivity without broadly suppressing protective humoral immunity.

## LIMITATIONS OF THE STUDY

Due to their low abundance, we were unable to identify T_FR_ in the single-cell sequencing data from the R848 treatment cohort, precluding trajectory and cell-cell interaction analyses of this population in the global dataset. Single-cell analyses of sorted populations nevertheless enabled detailed characterization of transcriptional states and repertoire features in isolation. Together with intravital tracking and functional perturbation experiments targeting T_FR_, these data allowed us to infer key aspects of T_FR_ identity and function in response to inflammation. However, we did not directly assess TCR specificity or antigen reactivity of T_FR_ in this model, because such analyses remain challenging given the intrinsic diversity of the response. Finally, analyses in SLE patients were restricted to peripheral blood and relied on correlative associations, as lymphoid and tissue-resident compartments could not be accessed.

## MATERIALS AND METHODS

### Transgenic mouse models

B6.129(Cg)-*FoxP3^tm4(YFP/icre)Ayr^*/J ^53^ and B6.129S(FVB)-*Bcl6^tm1.1Dent^*/J ^54^ were purchased from Jackson laboratories (JAX stock no. 016959 and no. 023727, respectively) and intercrossed. FoxP3-YFP-iCre Bcl6^flx/flx^ mice (T_FR_-deficient mice) used in experiments were Cre homozygous (females) or hemizygous (males) and Bcl6^flx^ homozygous. Cre negative Bcl6^flx/flx^ mice and C57BL/6J mice were used as controls. FoxP3-DTR/GFP mice (B6.129(Cg)-*Foxp3^tm3(Hbegf/GFP)Ayr^*/J) (JAX strain no. 016958) ^38^ were used to visualize T_REG_ and T_FR_ cells. FoxP3-DTR/GFP mice express FoxP3 bicistronically with human diphtheria toxin receptor (DTR) fused to a green fluorescent protein (GFP). Homozygous females and hemizygous males were used in experiments. 564Igi mice (B6.Cg-^Ightm1(Igh564)Tik^Igk^tm1(Igk564)Tik^/J) (JAX stock no. 032723) ^35^ were originally provided by Theresa Imanishi-Kari and were maintained in-house. 564Igi mice served as a positive control for autoreactivity in mass spectrometry experiments. Mice were bred and maintained under specific pathogen-free (SPF) conditions in individually ventilated cages under regulated and ambient temperature (20-22 °C) and humidity and kept on a 12-hour light/dark cycle with standard chow and water *ad libitum*. All experimental procedures involving animals in this study were approved by local authorities (Animal Experiments Inspectorate, Denmark, protocol number 2022-15-0201-01288).

In this study, some previously published murine datasets were reanalyzed ^55^. Flow data from Cre-negative Bcl6^flx/flx^ mice were previously included as Cre-negative controls for Aicda-Cre-positive mice ^55^. These experiments were run in parallel with the FoxP3-Cre-positive mice presented in this manuscript. The shared use of control mice across two independent experiments, when run in parallel, is consistent with the 3R principles. Intravital imaging datasets were furthermore used for method validation ^44^.

### Human PBMC samples

PBMC samples from 16 SLE patients were part of the Aarhus SLE biobank and were originally obtained at the Department of Rheumatology, Aarhus University Hospital. Inclusion criteria for the biobank were fulfilment of the 1997 American College of Rheumatology (ACR) classification criteria for SLE ^56^, age 18 or above, understanding and speaking Danish. Exclusion criteria were infection, ongoing cancer treatment, and incapacitation. After informed written consent, clinical data, including the SLE Disease Activity Index (SLEDAI) ^57^, were collected. Blood samples for research purposes were drawn at the same time as samples for routine biochemical assessments. Control samples (n = 7) were obtained from blood donors at the blood bank of Aarhus and Aalborg University Hospitals, Denmark. The Danish Data Protection Agency and the Regional Committee on Health Research Ethics approved the study (case no. 1-10-72-288-18). Blood was collected after informed consent according to the Declaration of Helsinki.

### Resiquimod (R848) treatment

Epicutaneous R848 application was carried out as described previously ^55^, 3 times/week for 2, 3, 4, 10, or 12 weeks. High-dose treatment involved 4 strokes of the R848-soaked cotton applicator, which, for the low-dose treatment, was reduced to 2 strokes. Mice were either euthanized immediately at the end of the treatment period or left untreated for 2-6 weeks before euthanasia. Detailed information on experimental timelines can be found in respective figures. Data from untreated control mice are indicated as untreated (UT) or as week 0.

### Murine tissue collection

Mice were euthanized at different time points as indicated in figures. Immediately before euthanasia, mouse body weight was documented. Blood was collected by decapitation under anesthesia with isoflurane for flow cytometry analysis and serum isolation. For flow cytometry, blood was collected in 200 µl PBS with 5 mM EDTA. For serum extraction, blood was left to coagulate for 30 min at room temperature. Serum samples were then centrifuged twice; first at 3,000 *g*, then at 20,000 *g*. Serum was stored at –80 °C until analysis. Lymph nodes were removed, cleaned of fat tissue, and placed into ice-cold FACS buffer (PBS, 2% heat-inactivated fetal bovine serum (FBS), 1 mM ethylenediaminetetraacetic acid (EDTA)). Spleens were removed, weighed on a precision scale (Mettler Toledo MS303S), and either placed into ice-cold FACS buffer for flow cytometry, freshly cut using a vibratome for two-photon imaging, or embedded into cryomolds and frozen down in OCT medium (TissueTek). Kidneys were removed, cut in half, and then fixed in 4% w/v paraformaldehyde (PFA) for 2 nights at 4 °C, afterwards washed in 10 mM PBS several times, dehydrated in 70%, 96%, and 99% ethanol for 2 hours, respectively, before they were transferred to xylene overnight. The day after, kidneys were embedded in paraffin.

### Immunofluorescence microscopy of spleen tissue

A Cryostar NX70 Cryostat (ThermoFisher) was used to cut 20 µm thick spleen. Sections were mounted on SuperFrost+ glass slides (Epredia, REF J1800AMNZ) and were acetone fixed (Merck, catalogue number 1000141000). Briefly, the spleen samples were rinsed in PBS and fixed in acetone for 10 minutes at room temperature. Thereafter, the slides were washed twice with PBS 0.1% w/v sodium azide. Biotinylated antibodies were diluted in staining buffer (PBS, 2% v/v FBS, 0.1% w/v sodium azide and centrifuged for 10 min at 10,000 *g* at 4 ℃ to remove antibody aggregates. Slides were stained for 4 hours at 4 ℃ in the dark, then washed once with staining buffer for 5 minutes and three times with PBS 0.01% v/v Tween-20 (Merck, product no.: 8.17072). Then, primary conjugated antibodies and conjugated streptavidin were diluted in staining buffer (PBS, 2% v/v FBS, 0.1% w/v sodium azide and centrifuged for 10 min at 10,000 *g* at 4 ℃ to remove antibody aggregates. Slides were then stained overnight at 4 ℃ in darkness. Slides were washed once with staining buffer, and three times for 5 minutes with PBS 0.01% v/v Tween-20, spot-dried and mounted using Fluorescence Mounting Medium (S3023, Dako). After drying, slides were sealed airtight with nail polish and stored in the dark at 4 ℃ until imaged. Imaging was done using a Z 22 Olympus VS120 Upright Widefield fluorescence slide scanner equipped with a digital monochrome camera (Hamatsu ORCA Flash4.0V2) and a 2/3” CCD camera, as well as single band exciters and a filter wheel with single-band emitters (DAPI, FITC, Cy3, Cy5, and Cy7). Fiji v. 2.1.0/1.53c was used for image processing and analysis. For analysis, FoxP3^+^ cells were counted within IgD exclusion zones and normalized to the measured GC area. The following antibodies were used: Ki67-eFluor660 (1/500, 50-5698-82, ThermoFisher Scientific), FoxP3-PE (1/500, 560408; BD Bioscience), CD4-Biotin and Streptavidin-AF488 (1/500, 553045, BD Biosciences and S32354, ThermoFisherScientific), IgD-PacificBlue (1/250, Invitrogen, clone: ALY7, 48-0443-82).

### Immunohistochemistry (IHC) and PAS histochemistry of kidney tissue

Formalin-fixed paraffin-embedded kidney tissue was sectioned at 4 µm on a microtome (Microm HM355S) and dried at 60°C for 1 h. Tissue sections were dewaxed in xylene, hydrated in graded alcohols and rinsed in deionised water. Antigen retrieval and peroxidase blocking were performed using EnVision FLEX+ reagent (K8005 and SM801 DAKO, respectively). Tissue sections were then incubated in 0.2% BSA (Bovine Serum Albumin) blocking buffer solution for 30 min. Primary antibodies (goat anti-mouse IgA, 1040-01-Southern Biotech [0.02 mg/ml] / goat anti-mouse IgG2c, 1079-08-Southern Biotech [0.02 mg/ml] / goat anti-mouse IgG (H+L), A160800-Invitrogen [0.0025 mg/ml]) were diluted in blocking buffer solution and incubated overnight at 4 °C. A secondary HRP-linked anti-goat antibody (AP180P-Sigma-Aldrich [0.0008 mg/ml]) was applied for 60 min at room temperature. The chromogenic detection was performed with EnVision FLEX DAB+/Substrate Buffer (DAKO), followed by hematoxylin counterstaining, dehydration and mounting.

Immunohistochemical stains were scored semi quantitatively as High and Low intensity stain per individual glomerulus. On each IHC stained slide 50 glomeruli were scored for antibody deposition and the absolute number of High– and Low– intensity-stained glomeruli was recorded.

Periodic Acid Schiff (PAS) histochemical stain was performed with PAS kit (RRSK15, Atom Scientific) following the manufacturer’s instructions. In brief, 4 µm thick FFPE mouse kidney tissue sections were dewaxed in xylene, hydrated through descending alcohol grades and rinsed with deionised water. The sections were oxidized with periodic acid, rinsed, stained with Schiff reagent, rinsed again, counterstained with hematoxylin, differentiated in 0.5% acid alcohol, rinsed with tap water and finally dehydrated and mounted on glass slides.

Glomerular lesions were graded semi quantitatively on a scale of 0 to 2+ for mesangioproliferation, endocapillary proliferation, mesangial matrix expansion, capillary collapse, hyalinosis, sclerosis and fibrosis (Score 0: <10%; Score 1:10–50%; Score 2: >50% of the glomeruli examined) on H&E and PAS-stained slides. Scores were calculated for each mouse based on at least 50 glomeruli.

### Anti-dsDNA antibody measurements by time-resolved immunofluorometric assay (TRIFMA)

FluoroNunc Maxisorp 96-well plates were coated with 100 µg/mL salmon sperm dsDNA (Invitrogen AM9680) in PBS and incubated overnight at 4°C. Wells were blocked with 200 µl TBS containing 1% w/v bovine serum albumin (BSA) for 1 hour at room temperature, then washed 3 times with TBS containing 0.05% v/v Tween-20 (TBS/Tw). Samples, standards, and quality controls were diluted in TBS/Tw containing 5 mM EDTA and 0.1% w/v BSA and were loaded onto the plate in duplicates. After overnight incubation at 4 °C, wells were washed 3 times with TBS/Tw and incubated for 1 hour at room temperature with biotinylated goat anti-mouse IgG2c (Southern Biotech, Cat.: 1077–08, diluted in TBS/Tw to 1 μg/ml). Wells were washed 3 times in TBS/Tw. Then Eu^3+^-labeled streptavidin diluted 1:1000 in TBS/Tw containing 25 µM EDTA was added to the wells and incubated for 1 hour at room temperature. Finally, the wells were washed 3 times in TBS/Tw, and 200 µL enhancement buffer was added. The plate was shaken for 5 minutes, and then the counts were read using a time-resolved fluorometry plate reader (Victor X5, PerkinElmer).

### Total IgG2c measurements

FluoroNunc Maxisorp 96-well microtiter plates were coated with goat-anti-mouse IgG2c antibody (Southern Biotech, Cat.: 1077–01, diluted to 1 µg/ml in PBS) overnight at 4 °C. Wells were blocked with human serum albumin (CSL Behring, Cat.: 109697), 1 mg/ml in TBS (Fisher Scientific, Cat.: BP2471–500) containing 0.09% w/v sodium azide, for 1 h at room temperature, and subsequently washed three times with TBS/Tw. Serum samples, standards and internal controls were resuspended on a vortex mixer and diluted in TBS/Tw containing 100 μg/ml of heat-aggregated human Ig (hIg Gammanorm, Octapharma, Cat.: 478393) and added to wells; each dilution was added in duplicate wells, incubated overnight at 4 °C or 2h at room temperature, and subsequently washed three times with TBS/Tw. Wells were next incubated for 2 h at room temperature with biotinylated goat anti-mouse IgG2c (Southern Biotech, Cat.: 1077–08, diluted in TBS/Tw to 1 μg/ml). Wells were subsequently washed three times with TBS/Tw. Next, wells were incubated 1 h at room temperature with Eu^3+^-labelled streptavidin (PerkinElmer, Cat.: 1244–360) diluted 1:1000 in TBS/Tw containing 25 μM EDTA and subsequently washed three times with TBS/Tw. Enhancement buffer was added to wells, which were then shaken for approx. 5 min on a plate shaker, then analyzed on a Victor X5 Multilabel Plate Reader (PerkinElmer). Mouse IgG2c (Southern Biotech, 5300-01B) was used as standard.

### Flow cytometry analysis of murine tissue

Spleen pieces and lymph nodes were mechanically macerated in 1.5 mL tubes containing 500 µl ice-cold FACS buffer (PBS, 2% heat-inactivated FBS, 1 mM EDTA) using plastic pestles and then filtered through 100 µm cell strainers. Spleen samples were additionally incubated with red blood cell (RBC) lysis buffer (155 mM NH_4_Cl, 12 mM NaHCO_3_, and 0.1 mM EDTA) at room temperature for 5 min, pelleted, then resuspended in 750 µl FACS buffer. Peripheral blood mononuclear cells (PBMCs) were isolated from blood samples using Lympholyte M (Cedarlane Labs), washed, and finally resuspended in 100 µl FACS buffer. One hundred µl of each cell suspension was plated with 20 µl Fc-block solution and incubated for 5 min on ice. Antibodies and a viability dye were diluted in FACS and brilliant stain buffer (BD Horizon™) (50:50) and 100 µl of this antibody mix was added to each sample and incubated for 30 min on ice. After washing, cells were fixed for 30 min with 0.9% PFA in PBS, then resuspended in 200 µl FACS buffer. For CXCR5 staining, samples were incubated at 37°C with anti-CXCR5-biotin for 30 min, then washed, before regular staining on ice with streptavidin-BV421 and other cell-surface markers. For staining of intracellular targets (FoxP3, Helios, anti-GFP), cells were fixed and permeabilized after cell-surface marker staining using the eBioscience Foxp3/Transcription Factor Staining Buffer Set (00-5523-00, ThermoFisher), before incubation with antibodies targeting intracellular targets. Flow cytometry evaluation was performed using a 4-laser (405 nm, 488 nm, 561 nm, 640 nm) BD LSR Fortessa flow cytometer with 16 fluorescence detectors (Becton Dickinson, San Jose, CA) or a 4-laser (405 nm, 488 nm, 561 nm, 637 nm) NovoCyte Quanteon 4025 flow cytometer with 25 fluorescence detectors (Agilent, Santa Clara, CA). Data was analysed using FlowJo versions 10.8.1-10.10.0 software (Tree Star, Inc., BD Life Sciences). The universal gating strategy for each dataset is shown in the respective supplementary figures.

### Autoantibody screening on HEK293TN cells

Round glass coverslips (Thermo Fisher Scientific, Menzel Gläser, Ø 18 mm, #1 thickness, Lot# 0584) were UV-light-treated for 20 minutes on each side and transferred to 12-well plates (Sarstedt, Cat# 83.3921). 30,000 HEK293TN cells were seeded per well and grown for 24 h in 2 mL culture medium (High-Glucose Dulbecco’s Modified Eagle Medium (DMEM) (Gibco, Cat# 11965-092) supplemented with 10% heat-inactivated FBS (Sigma Aldrich, Cat# F7524) and 1% v/v penicillin-streptomycin (P/S) (Thermo Fisher Scientific, Cat# 15140-122)). Culture medium was sterile filtered before use. Cultures were maintained at 37 °C in a humidified incubator with 5% CO_2_ (Thermo Fisher Scientific, Series 8000DH CO_2_ Incubator). After 24 hrs, cells were washed in PBS and fixed with formalin (Sigma Aldrich, Cat# F1635) diluted 1/10 in PBS (700 µl/well) for a 30 min incubation on an orbital shaker (Grant, PSU-10i, 70 rpm) at room temperature. Cells were then washed in PBS and incubated with TBS for 15 min, followed by another wash in PBS. For permeabilization, cells were incubated with 0.1% Triton X-100 (Sigma-Aldrich, Cat# X100) in PBS containing 2% FBS (Sigma-Aldrich, Cat# F7524) for 15 min on an orbital shaker at RT (700 µl/well). Afterwards, they were washed in PBS. Cells were then incubated overnight with mouse serum from 4 different cohorts diluted 1:160 in PBS containing 2% fetal bovine serum (FBS, Sigma-Aldrich, Cat# F7524) in a humid chamber at room temperature on an orbital shaker (Grant, PSU-10i, 70 rpm). The following day, cells were washed twice in PBS containing 2% FBS and incubated with biotinylated goat anti-mouse IgG2c (1:600, Cat# 1077-08, SouthernBiotech) for 4 h at room temperature on an orbital shaker. After incubation, cells were washed twice in PBS with 2% FBS and incubated with AlexaFluor488-conjugated streptavidin (1:1000 S32354 Thermo Fisher Scientific) and DAPI (1:2000, 422801 Nordic Biosite) for 2 h in the dark at room temperature on an orbital shaker. After incubation, cells were washed twice in PBS with 2% FBS again and the coverslips with the adherent cells were mounted on microscope slides (Thermo Fisher Scientific, Epredia™ J1800AMNZ) with 12 µl DAKO fluorescence mounting medium (Agilent Technologies, Cat# S3023) and stored for at least 24 h at 4°C before imaging. Slides were imaged using a Zeiss LSM800 laser-scanning confocal microscope. A 40x oil immersion objective with a numerical aperture of 1.4 was used. Single-plane images of four fields of view with non-dividing cells and non-overlapping nuclei were taken from each sample. The image setup was 1024×1024 px, a scan speed of 4.12 µs, bidirectional imaging, 4x averaging repeated per frame based on mean intensity, and an optical section of 0.4 µm across all channels.

### Autoantigen pull-downs for autoreactivity screening by mass spectrometry

The protocol was based on our earlier work ^8^. Briefly, four female C57BL/6JRj mice aged 6-8 weeks (Janvier labs) were euthanized by cervical dislocation. The spleens were extracted, and each was placed into 600 µl ice-cold buffer (20 mM imidazole, 10% w/v sucrose, 2 mM EDTA, pH 7.4) containing protease inhibitor cocktail (Thermo Fisher 78425), then homogenized in a bullet blender (Next Advance) at speed 8 for 30 seconds. The homogenate was centrifuged at 3,000 *g* for 10 min, at 4 °C, and the supernatant was aliquoted and frozen at −20 °C until use. For pre-clearing of supernatants, sepharose beads with Protein G (Cytiva 17088601 Gammabind Sepharose) were washed in Buffer 3 (0.2 M glycine, pH 2.5), then equilibrated in Buffer 1 (PBS containing 0.05% Tween-20, 0.1% w/v BSA, 0.1% NaN_3_ and 2 mM EDTA). 350 µl beads were incubated with 350 µl pooled spleen supernatant and 350 µl Buffer 1, for 30 min, end-over-end at 4 °C. Beads were pelleted at 100 *g* for 3 min at 4 °C, and the supernatant was recovered. For each sample, 10 µl of serum was incubated with 50 µl of the pre-cleared supernatant for 1 h on ice, then each was added 25 µl freshly washed and equilibrated Sepharose beads and incubated another 30 min end-over-end at 4 °C. Beads were recovered by centrifugation at 100 *g*, for 3 min at 4 °C, then washed 4 times with 500 µl Buffer 2 (PBS containing 0.05% v/v Tween-20, 0.1% NaN_3_, and 2 mM EDTA). After the washes, bound proteins were eluted with 55 µl Buffer 3, and the eluate was neutralized with 13 µl 1 M Tris-HCl, pH 8.0. Eluates were frozen at −20 °C until MS analysis.

### LC-MS-based proteomic identification of autoantigens

The label-free proteomics analyses were performed similarly to a previous protocol ^58^. Briefly, 25 µl eluate was mixed with SDS-PAGE sample buffer, and subjected to SDS-PAGE followed by reduction, blocking of reduced cysteine residues, and in-gel trypsin digestion. Eluted peptides were purified on PepClean C18 Spin Columns and vacuum dried. Peptides were then separated by liquid chromatography (Easy-nLC 1200, Thermo Scientific) using a pre-column (Acclaim PepMap 100, 75 µm × 2 cm, Nanoviper, Thermo Scientific) and an analytical column (EASY-Spray column, PepMap RSLC C18, 2 µm, 100 Å, 75 µm × 25 cm) in a 60 min gradient of 4–40% acetonitrile in 0.1% formic acid, coupled to the mass spectrometer Q-Exactive HF-X Hybrid Quadrupole Orbitrap (Thermo Scientific, Bremen). Full-scan (MS1) resolution was set to 60,000 (at 200 m/z) and the scan range was between 372 and 1800 m/z. Up to the 12 most intense peaks in MS1 were fragmented in MS2 using data-dependent acquisition. MS2 resolution was set at 15,000. Unassigned and +1 charge states were excluded from fragmentation, and a dynamic exclusion of 15 s was used. Proteins were identified and quantified using the Andromeda algorithm in MaxQuant (version 1.5.3.30) ^59^ to search against a mouse sequence database (*Mus musculus* proteome with 17,544 reviewed sequences downloaded from Uniprot database, https://www.uniprot.org/ on 04/12/2022). The “match between runs” was applied, and iBAQ data, were used for quantification. Data were filtered to exclude IgH, IgK, and IgL chains derived from the pull-down antibodies, as well as keratins stemming from human contamination. iBAQ values were log2(x+1) transformed before volcano plotting. P values from unpaired t tests were adjusted to q values using the two-stage linear step-up procedure of Benjamini, Krieger, and Yekutieli (FDR = 5%). Additionally, raw iBAQ values were imported into R (v. 4.5.0), and then scaled using the scale function. Any columns with missing data were removed. Principal components were calculated using the prcomp function of FactoMineR (v. 2.8), UMAP was performed using uwot (v. 0.2.3), and plots were rendered using ggplot2 (v. 3.5.2).

### Single-cell sequencing of the R848 treatment cohort

Single-cell sequencing data for the R848 treatment cohort were based on two independent experiments. The first experiment included a spleen sample and both IngLNs from 4 mice, one in each of the four groups. The 4 spleen samples and the 4 IngLN samples were barcoded and loaded onto each their 10x Chip K Chromium Lane. The second experiment included a spleen sample from 8 additional mice, two mice from each of the four groups. Samples were barcoded and two pools were generated, each containing one spleen sample from each group. The resulting two pools were loaded into each their respective 10x Chromium Chip K lanes. Libraries were prepared according to standard 10x Genomics protocols and quality-controlled using Qubit and TapeStation. The first experiment was sequenced by BGI Europe, Copenhagen, on a DNBSEQ-G400 platform with 26-bp and 90-bp read lengths for read 1 and read 2, respectively. The second experiment was sequenced at the NGS Core at MOMA, Aarhus University Hospital, on an Illumina NovaSeq X Plus using a 25B 300-cycle flow cell with a 150–10–10–150 bp read structure, including approximately 25% PhiX spike-in. Sequencing data were preprocessed with CellRanger and then underwent a standard QC pipeline in Seurat v. 5.3.0 ^60^ using R version 4.5.0. Additional packages included ggplot2 (v. 3.5.2), ggrepel (v. 0.9.6), sctransform (v. 0.4.2), magrittr (v. 2.0.3), dplyr (v. 1.1.4), assertthat (v. 0.2.1), gprofiler2 (v. 0.2.3), Cairo (v. 1.6.2), ggpubr (v. 0.6.3), decoupleR (v. 2.14.0), dorothea (v. 1.20.0), tidyr (v. 1.3.2), tibble (v. 3.3.1), stringr (1.6.0), and clustree (v. 0.5.1). Global clustering identified 20 clusters covering all hematopoietically derived splenocyte subsets. Five clusters expressing Cd79a, Ms4a1, and/or Sdc1, were identified as B-lineage and extracted for B/plasma-cell subclustering. Similarly, 6 clusters expressing Cd3e, Cd4 and/or Cd8a were identified as T-lineage and extracted for T cell subclustering. V(D)J sequence analysis was performed using scRepertoire v. 2.5.2 ^61^. QuietVDJgenes function was modified from the original code provided by Nick Borcherding et al., to accommodate *Mus musculus* genes.

### Surgical preparation for serial intravital microscopy of the spleen

To allow serial intravital imaging of splenic GCs for 2 weeks, a 14 mm diameter wide biocompatible custom-made AIW, consisting of a rimmed titanium ring (SOLIDPART Denmark) and a passivated (Poly-2-methyl-2-oxazoline (PMOXA) glass coverslip, was implanted into the abdominal wall over the spleen 24-72 h before the first imaging day. This was based on a protocol developed by us ^44^, using an adapted AIW ^62^ to specifically enable serial intravital imaging of abdominal organs using an upright two-photon microscope. The day before AIW implantation, hair on the left flank of the animals was removed. Mice were anesthetized with isoflurane in medical air and 50% oxygen (induction: 3.5% isoflurane; maintenance 1.2–1.8% isoflurane; flow rate: 0.6-1.2 L/min, Anesthetic Vaporizer UNO300VAP). The mice received analgesia with buprenorphine in 0.9% NaCl (0.1 mg/kg bodyweight i.p.) 15 min prior to surgery, application of protective eye ointment (Visc-ophtal eye gel 2 mg/g, Orifarm) and skin disinfection with chlorhexidine. The AIW was implanted into the abdominal wall of the left flank as previously described ^44^. Immediately after surgery, mice received meloxicam (1 mg/kg bodyweight s.c.). Postoperative analgesia (0.009 mg/ml buprenorphine in drinking water and 1 mg/kg bodyweight meloxicam s.c./day) was carried out for 2 further days after surgery day (mice were adapted to the taste of analgesic water 2 days prior to surgery). We performed daily wound checks to ensure appropriate wound healing, and to support recovery, mice were provided with buprenorphine-soaked chow for the first 3 postoperative days and with easily accessible dry chow on the cage bottom throughout the entire imaging period. Mice were single housed after surgery to avoid damage to the AIW. A detailed methodological description can be found elsewhere ^44^.

### Serial intravital two-photon microscopy

Mice were imaged 5 times over 15 days to track longitudinal changes in GCs. 24 h before each imaging time point, mice received 5 µl αCD35-iFluor647 (0.43 mg/ml) in 95 µl sterile PBS i.v. to label FDCs and mark the light zone of GCs. αCD35 was conjugated to iFluor647 using an in-house labeling protocol to a labeling density of 12. For intravital imaging, mice were anesthetized with isoflurane in medical air enriched with 33% oxygen (induction: 3.5% isoflurane; maintenance 0.7-1.5% isoflurane; flow rate: 0.6-1.2 L/min, using either SomnoSuite apparatus (Kent Scientific, United States) or Anesthetic Vaporizer UNO300VAP). Corneas were protected with eye ointment, and mice were hydrated with 2x 100 µl 0.9% NaCl s.c. before and after imaging. Mice were carefully mounted in a custom 3D-printed holder for upright intravital imaging ^44^ and placed on a servo-controlled heating plate. Microscopy was carried out with an Olympus FVMPE-RS multiphoton system with Fluoview FV31S software, equipped with an integrated x25 water immersion objective with an isolated tip and numerical aperture of 1.05 (Olympus, XLPLN25xWMP2: WD 2.00 mm), a MaiTai DeepSee Olympus laser (Spectra Physics), and 2 high-performance multi-alkali PMT and 2 GaAsP PMT detectors. Images were acquired with two sequential excitation wavelengths, λ _Ex_ = 840 nm and λ _Ex_ = 940 nm, to capture distinct spectral features: λ _Ex_ = 840 nm was used to excite iFlour647 (CD35), and λ _Ex_ = 940 nm was used to excite GFP (FoxP3) and observe SHG (collagen). Fluorescence emission was collected in 4 channels (to balance signal to autofluorescence) in both excitation tracks using the following filter sets: Ch 1 & 2: 650/60 & 573/75, separated by an Olympus FV30-SDM570 primary emission beam splitter from Ch 3 & 4 (GaAsP): 520/30 & 452/45. An IR short pass filter, Olympus BA685RXD, was used to filter out emission light above 685 nm.

At the beginning of every imaging session, a 3D view of the spleen within the window’s field of view was acquired to compile an overview map, facilitating the reidentification of GCs over time. For more information, please refer to ^44^. Individual GCs were acquired in volumetric image stacks with a 5 µm step size, reaching a total depth of up to 220 µm, measured from the capsule to the deepest imaging point. At λ_Ex_ = 840 nm, images were acquired in galvo scanning mode without averaging, using 2 µs dwell time/pixel at 800 x 800-pixel resolution and at λ_Ex_ = 940 nm with 3 x line averaging, using 4 µs dwell time per pixel at 800 x 800-pixel resolution. Detector gains were adjusted at the beginning of each serial imaging cohort to balance the fluorescence emission of interest and autofluorescence and were then kept unchanged throughout each experiment. Laser power was increased exponentially in depth and adjusted to avoid signal saturation, phototoxicity, and bleaching while still detecting all fluorophores of interest throughout the entire 3D stack.

### Intravital image processing and image analysis

Images were processed and analyzed using Fiji (v2.14.0/154f) ^63^. Initially, 3D imaging stacks from all imaging days for each GC were standardized in stack position and stack size by manually matching landmarks of the capsule, vasculature, and splenic trabeculae. Each longitudinal imaging set was finally visualized with the Multi Stack Montage plugin from the PTBIOP Update Site ^64^ to ensure consistency in stack standardization. For 3-dimensional visualization of the FDC network over time, an FDC network of each imaging day was reconstructed using the ‘Big Structure Detection in 3D’ module in Arivis Vision4D (v. 4.3.0, Arivis AG, Rostock, Germany) with manual thresholding applied on CD35-iFluor647 signal to account for signal variability and autofluorescence across serial intravital imaging days. Threshold accuracy was validated by direct comparison of segmented volumes with the corresponding raw micrographs, inspected plane-by-plane (Fig. S4A-C).

To quantify FDC network dynamics, the FDC network circumference was manually outlined in an average Z-projection using the built-in Z function in Fiji (25) on the λ Ex = 840 nm track (Fig. 3C and D, and Fig. 5C-F). Solely frames with positive CD35^+^ staining were selected for Z-projection to avoid overlay of red pulp macrophages above the FDC network and facilitate visualization. Before using the freehand tool to outline the FDC network, the “Despeckle” filter in Fiji was applied. Manual outline was eased by activating channel 1 (CD35^+^ and macrophages) and channel 3 while applying complementary pseudo colors (magenta and green) to both channels. Finally, the measured area of the circumference was normalized to the first observation timepoint to reveal relative size changes in the FDC network over time. For more information please refer to ^44^.

For T_FR_ quantification, both excitation tracks were merged to identify FoxP3-GFP^+^ cells (λ _Ex_ = 940 nm) in GCs (λ _Ex_ = 840 nm). GCs were identified based on their light zone (CD35-FDC labelling), and the entire GC area was determined based on TBM localization and absence of red pulp macrophages proximal to the FDC network. FoxP3^+^ cells were visualized based on their GFP expression ^38^ and T_FR_ cells were identified based on their localization in GCs, which included GFP^+^ cells localized within the CD35-labeled area, as well as those in very close proximity, especially close to TBMs. Before merging the iFluor647-channel (channel 1) from λ _Ex_ = 840 nm track and channels 2-4 from λ _Ex_ = 940 nm track, background illumination from the respective channels was individually corrected by rolling ball background subtraction (rolling ball radius 50 pixels). After merging “Despeckle” filter was applied. Lastly, the “Maximum 3D” filter was applied to further enhance the GFP-signal of single FoxP3^+^ cells (T_REG_ and T_FR_ cells). Finally, T_FR_ cells were counted manually by setting ROIs using the point tool throughout the entire GC volume, slice by slice. For more information on image processing to enhance the GFP signal, please see ^44^. For three-dimensional visualization of the entire GC and the spatial distribution of T_FR_ cells over time, a GC from each imaging day was reconstructed using the ‘Draw Object’ function in Arivis Vision4D (v. 4.3.0, Arivis AG, Rostock, Germany). GC outlines were determined by the area spared of red pulp autofluorescent macrophages in proximity to the FDC network. T_FR_ cell positions were determined from manually selected ROIs in Fiji and converted to sphere objects in Arivis Vision4D for visualization.

### Adoptive transfer experiment

Donor (FoxP3 reporter, FoxP3-DTR/GFP) and recipient (T_FR_-deficient, FoxP3-YFP-iCre Bcl6^flx/flx^) mice were R848-treated from Day 1 (3x/week). Spleens were harvested from donor mice on Day 3, and single-cell suspensions were prepared by mechanical dissociation followed by filtration through 100-μm strainers. Red blood cells were removed with RBC lysis buffer. To isolate CD4^+^FoxP3^+^ and CD4^+^FoxP3^−^ populations, negative magnetic selection was performed using magnetic-activated cell sorting (Miltenyi Biotec #130-091-051, Miltenyi Biotec 130-042-303). Total cells were incubated with lineage-depletion biotin antibody cocktail (NK1.1 (clone PK136) Biolegend #108704, CD8a (clone 53-6.7) Biolegend #100704, B220 (clone RA3-6) Biolegend #103204, TER-119 Biolegend #116204, CD11b (clone M1/70) Biolegend #101204, CD11c (clone N418) Biolegend #117304, Gr1/Ly6g/c (clone RB6-8C5) Biolegend #108404, CD14 (clone Sa14-2) Biolegend #123306) followed by anti-biotin microbeads (Miltenyi Biotec #130-090-485), then passed over LS columns (Miltenyi Biotec #130-042-401) on a magnetic separator. The flow-through fraction containing negatively selected CD4^+^ T cells was collected and stained with live-dead marker, anti-CD4-PE-CF594, and streptavidin-BV421, then subjected to FACS on a Bigfoot cell sorter equipped with six lasers (349 nm, 405 nm, 455 nm, 488 nm, 561 nm, and 640 nm) and 52 fluorescence detectors (Thermo Fisher, Fort Collins, CO). Cells were gated as live, CD4^+^ singlets and sorted into GFP⁺ and GFP⁻ fractions. Sorted populations were collected in sterile-filtered HBSS with 12.5% heat-inactivated murine serum. The GFP⁻ fraction was labeled with a Qtracker 655 Cell Labeling Kit (Thermo Fisher, Q25021MP) following the manufacturer’s instructions. QC was performed by flow cytometry as described in Flow Cytometry section above, staining with streptavidin-BV421 (BD #563259), CD19-BV510 (BD #562956), CD4-PE-CF594 (BD #562314), TCRb-PE-Cy5 (BD #553173), Viability dye eF780 Invitrogen #65-0865-14), and additionally reading out GFP and Qdot655 signals (see Fig. S6A). Thereafter, GFP^+^ and GFP^−^ populations were pooled and transferred into recipient mice. Each mouse received 3 x 10^5^ GFP^+^ cells and 1.3 x 10^6^ GFP^−^ cells in 200µl HBSS by retroorbital injection. R848 treatment continued until day 17. On day 16, the FDC-network was intravitally labelled with 2 µg anti-CD35-iFluor647 in 100 µl sterile PBS intravenously 24 h before euthanasia. CD35 (clone 8C12, BD Biosciences 558768), was conjugated to iFluor647 (AAT Bioquest, Cat#1031ABD-1031, Nordic Biosite) using an in-house labeling protocol (final concentration 0.43 mg/ml and labeling density 12). On Day 17, mice were euthanized for analysis. The marginal zone was labeled only for fresh spleen explant imaging and was visualized by injecting 5 µg CD169-PE (142404; BioLegend) diluted in 100 µl PBS i.v. 10 min before euthanasia. Spleens were harvested, and half of each spleen was freshly cut into 400-800 µm-thick slices using a vibrating blade microtome (Leica VT1200 S) for fresh two-photon imaging (as previously described ^44^). The other half of the spleen was processed into single-cell suspensions, and flow cytometric staining was performed as described in the Flow Cytometry section above (gating tree in Fig. S6B).

### two-photon imaging of freshly harvested and stained thick spleen sections

Fresh explant two-photon microscopy was performed on the same multiphoton system, using similar hardware settings to those used during intravital microscopy. Images were acquired with two sequential excitation wavelengths, λEx 840 nm and λEx 940 nm, to capture distinct spectral features: λEx 840 nm was used to excite iFluor647 and PE, and λEx 940 nm was used to excite GFP, PE, and observe SHG. 3D image stacks were acquired using the MATL tile scan function (30% overlap) in Fluoview (2×2 – 3×3 single tiles) to capture the entire white pulp area. Stack acquisition with 3 µm step size reaching a total depth of up to 115 µm (measuring from cut surface to deepest imaging point). All images were acquired in galvanometer scanning mode with 3x line averaging, 4 µs dwell time/pixel at 800 x 800 pixels resolution. Detector Gains were adjusted at the beginning of each experimental cohort to balance the fluorescent emission of interest and autofluorescence and then kept unchanged. Laser power was increased exponentially in depth to avoid signal saturation and bleaching while still illuminating all fluorophores of interest throughout the entire 3D stack.

After imaging, the slice orientation and approximate localization of the imaged area were photo documented. Then, spleen slices were carefully removed from the imaging chamber and placed in an Eppendorf tube with 4% PFA overnight at 4°C. After fixation, slices were transferred into a tube containing PBS and 0.1% sodium azide at 4°C until further permeabilization and staining (ca. 2 weeks of storage). To visualize both Helios^+^ cells and the PALS of the freshly imaged white pulp areas, thick spleen slices were stained with anti-CD3-AlexaFluor594 (BioLegend; Clone 17A2), and anti-Helios-AlexaFluor488 (Clone: 22F6, BD #563950). Prior to staining, spleen sections were washed in PBS (3 × 15 min) on a shaking table at room temperature in the dark. Samples were then incubated overnight in the dark at room temperature in blocking buffer (5% FBS and 0.5% Triton X-100 in PBS with 0.1% sodium azide) on a shaker. The following day, the blocking solution was replaced with a staining buffer (2.5% FBS and 0.5% Triton X-100 in PBS with 0.1% sodium azide) containing CD3-AlexaFluor594 (1:70), and slices were incubated for 48 h on a shaking table at room temperature in the dark. After 48 h incubation, anti-Helios-AlexaFluor488 (1:100) was added directly to the same staining solution without intermediate washing, and samples were incubated for an additional 48 h. The FoxP3-GFP signal disappeared during staining and was only visible in fresh spleen explants. Intravital labeling of CD35-iFluor647 and CD169-PE largely withstood the fixation and staining process and supported reidentification of imaged areas from fresh explants.

Images of stained spleen sections were acquired at λEx 840 nm or λEx 940 nm excitation. Large 3D image stacks were acquired using the MATL tile scan function (30% overlap) in Fluoview in galvanometer scanning mode with 3-line averaging using 4 µs dwell time/pixel at 800 x 800 pixels resolution and stitched immediately post image acquisition. Detector gains were adjusted at the beginning of each experimental cohort to balance the fluorescence emission of interest and autofluorescence and were then kept unchanged. Laser power was adjusted to avoid signal saturation and bleaching while still illuminating all fluorophores of interest throughout the entire 3D volume. Laser power was increased exponentially with depth.

All representative images underwent image processing as described in the figure legends. Additionally, representative images from explanted spleens both fresh and stained, were registered to one another using the BigWarp plugin (97) in FIJI for manual landmark-based image alignment (thin-plate spline model). Autofluorescent macrophages (red pulp macrophages and TBMs) and CD169-PE labeling served as landmarks in all tracks.

### Single-cell gene expression and TCR repertoire analysis of sorted T cells

For the gene expression and TCR repertoire analysis of sorted T cells, spleen tissue from 4 R848-treated FoxP3-DTR-GFP mice was processed for FACS as indicated above using the following antibodies: B220-BV510, CD4-Qdot605, CD138-BV650, CD25-BV786, FoxP3-GFP, CD8-RB744, CXCR5-biotin+Streptavidin-PE, TCRb-PE-Cy5, CD38-PE-Cy7, CD95-A647, and Viability-eF780. T_REG_ cells, T_FR_ cells and T_FH_ cells were subsequently sorted based on GFP and CXCR5 expression using a Bigfoot cell sorter equipped with six lasers (349 nm, 405 nm, 455 nm, 488 nm, 561 nm and 640 nm) and 52 fluorescence detectors (Thermo Fisher, Fort Collins, CO), see gating tree in Fig S7A. After sorting, samples were barcoded and loaded on a single 10x Chromium Chip K Lane. The single-cell processing and analysis pipeline was similar to that of the R848 treatment cohort described above, but also used ggraph (v. 2.2.1), circlize (v. 0.4.16) and scales (v. 1.4.6).

### PBMC isolation and flow cytometry

PBMCs were isolated using CPT tubes (BD Diagnostics Vacutainers). Samples were centrifuged at 1800 *g* for 30 min at room temperature. Following centrifugation, the mononuclear cell layer was collected and transferred to 15 ml conical tubes. Following two washes with PBS, the cell pellets were resuspended in 10% DMSO with FBS and stored at −135 °C until use. All samples were thawed on the same day in a 37°C warm water bath. After thawing, cells were transferred and resuspended in 37 °C PBS containing 20% FBS. After centrifugation at 200 *g* for 7 min at room temperature, the supernatant was removed and the cells were resuspended in ice-cold FACS buffer. Cells were counted (Cellometer K2, Nexcelom Bioscience and ViaStain AOPI Staining Solution CS2-0106) and 1-3 x 10^6^ cells were plated per well. The staining procedures were similar to those for murine cells, and flow cytometric analysis was performed using a 4-laser (405 nm, 488 nm, 561 nm, 637 nm) NovoCyte Quanteon 4025 flow cytometer with 25 fluorescence detectors (Agilent, Santa Clara, CA). Data was analyzed using FlowJo version 10.10.0 (BD Life Sciences). The gating strategy is shown in Figure S8.

### Quantification and Statistical Analysis

Statistical analyses were performed using GraphPad Prism (v. 10.1.1 (270)). Values are expressed as means ± SD. P-values < 0.05 were considered significant. Statistical significance was determined using two-tailed paired t-tests; for grouped analyses, parametric or nonparametric tests were used, depending on normality and homogeneity of variances; and for multiple comparisons, one-way ANOVA, two-way ANOVA, or the Kruskal-Wallis test was used, with the appropriate post hoc tests. The Shapiro-Wilk test, Kolmogorov-Smirnov test, and QQ plots were used to assess the likelihood of normal or lognormal distributions. If data were more likely lognormally distributed, log-transformed was carried out to compute statistical significance and calculated p-values stem from log-transformed datasets. Data displayed in graphs are from non-transformed datasets unless stated otherwise. No outliers were excluded, but several datapoints were excluded for technical or biological reasons. Excluded samples, the n values, necessary data transformation, and the statistical tests used are indicated in respective figure legends.

## Acknowledgements

We thank the Laboratory Animal Facility, the Bioimaging Core Facility, and the FACS Core at Aarhus University for excellent technical assistance and support. We thank the technicians at the Department of Biomedicine for assistance with custom 3D-printing.

The 6-laser Bigfoot cell sorter is a generous gift from the Carlsberg Foundation, grant number CF21-0363.

## Author contributions

Conceptualization: LP, LV, SED

Methodology: LP, LV, YL, JP, LL, SED

Formal analysis: LP, LL, SED

Investigation: LP, LV, GW, SSJ, KG, YCLW, JKD, CFH, TRW, CS, MKP, JVH, KSK, SVF, JS, AS, AJH, LJ, AP, ADP, JP, LL, SED

Resources: SFG, AT

Data Curation: LP, JP, LL, SED

Writing—original draft: LP, SED

Writing—review & editing: All authors

Visualization: LP, LV, SED

Supervision: SFG, DS, SED

Project administration: SED

Funding acquisition: SED

## Data and materials availability

Processed data and raw data for single-cell RNA sequencing produced in this study are available via the Gene Expression Omnibus (GSE317196).

The mass spectrometry proteomics data have been deposited to the ProteomeXchange Consortium via the PRIDE ^65^ partner repository with the dataset identifier PXD073345.

Key reagents will be made available by the corresponding author upon reasonable request.

## Supplementary material

Figure S1-8 and full methods section are included with this manuscript.

